# Proteoliposomes as energy transferring nanomaterials: enhancing the spectral range of light-harvesting proteins using lipid-linked chromophores

**DOI:** 10.1101/609255

**Authors:** Ashley M. Hancock, Sophie A. Meredith, Simon D. A. Connell, Lars J. C. Jeuken, Peter G. Adams

## Abstract

Biology provides a suite of optically-active nanomaterials in the form of “light harvesting” protein-chlorophyll complexes, however, these have drawbacks including their limited spectral range. We report the generation of model lipid membranes (proteoliposomes) incorporating the photosynthetic protein Light-Harvesting Complex II (LHCII) and lipid-tethered Texas Red (TR) chromophores that act as a “bio-hybrid” energy transferring nanomaterial. The effective spectral range of the protein is enhanced due to highly efficient energy transfer from the TR chromophores (up to 94%), producing a marked increase in LHCII fluorescence (up to 3x). Our self-assembly procedure offers excellent modularity allowing the incorporation of a range of concentrations of energy donors (TR) and acceptors (LHCII), allowing the energy transfer efficiency (ETE) and LHCII fluorescence to be tuned as desired. Fluorescence Lifetime Imaging Microscopy (FLIM) provides single-proteoliposome-level quantification of ETE, revealing distributions within the population and proving that functionality is maintained on a surface. Our membrane-based system acts as a controllable light harvesting nanomaterial with potential applications as thin films in photo-active devices.

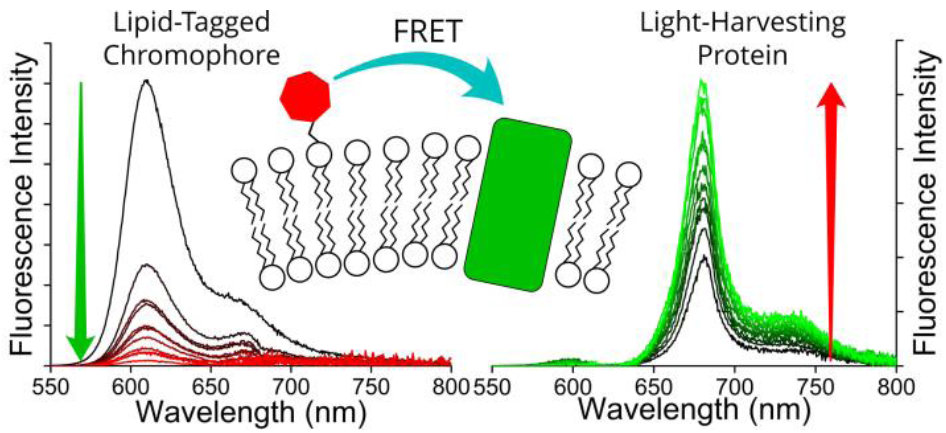
Table of Contents Figure.

## Main Text

Biological systems are a source of inspiration in solar energy research and nanotechnology,^1-6^ as the early stages of energy absorption and transfer in photosynthesis have an efficiency approaching unity.^7^ The membrane proteins which undertake photosynthetic light harvesting (LH) act in a conceptually similar way to a satellite dish, whereby antenna proteins channel energy towards Reaction Centre type proteins which perform photochemistry.^8^ LH membrane proteins have a 3-D polypeptide structure which acts as a scaffold to precisely arrange chromophores at high density in such a manner that energy absorption and transfer are extremely efficient.^9^ Light-harvesting complex II (LHCII)^10^ acts as the primary antenna protein for plants and is estimated to be the most abundant membrane protein on Earth due to its high density in the membranes of chloroplasts. LHCII has a heterotrimeric structure with each monomer containing fourteen chlorophyll and four carotenoids:^10^ this combination of chromophores gives the protein complex a high absorption coverage across the visible spectrum aside from the ‘green gap’ of minimal absorption between 520-620nm. Several studies have filled this green gap with complementary chromophores which absorb strongly in this spectral region and transfer energy to the protein via Förster Resonance Energy Transfer (FRET). This was first demonstrated using chromophores which were covalently attached to LH membrane proteins isolated in detergents, including synthetic organic compounds^11-14^ or quantum dots.^15, 16^ FRET has also been demonstrated between non-covalently coupled chromophores incorporated into lipid bilayers^17-19^ or polymer micelles.^20-22^ LHCII has previously been integrated into model membranes, e.g., lipid vesicles^23-28^ or lipid nanodiscs,^28, 29^ and these have been studied to understand its biophysical properties, but not to enhance its spectral range. Adding the chromophore to the lipid membrane would have several advantages compared to covalent modification of LH proteins: (i) lipids provide a more native environment for membrane proteins than detergent, (ii) membranes readily adsorb to hydrophilic solid supports so would be compatible with surface-based nanotechnologies, (iii) non-covalent systems allow greater flexibility to change the chromophore concentration (or type), (iv) membranes allow the potential to co-assemble other components to make for a modular system (e.g. other photosynthetic proteins or any small amphiphiles).

Solution-based fluorescence spectroscopy is typically used to assess FRET in ensemble measurements. Fluorescence Lifetime Imaging Microscopy (FLIM) has been previously applied to map the fluorescence of LHCII proteins within nanoscale 2-D clusters,^30^ micron-sized 3-D crystals^31^ and proteoliposomes,^25^ quantifying its fluorescence quenching (directly related to FRET) with ∼300 nm resolution. Here, we present a new proof-of-concept system where LHCII is non-covalently interfaced with additional chromophores in the lipid membrane to enhance its spectral range over the ‘green gap’. Samples are characterized with both ensemble spectroscopy and FLIM to provide a detailed quantitative assessment of their composition and FRET properties. Our results demonstrate the modularity, consistency and potential for highly efficient energy transfer of the non-covalent LHCII-chromophore systems.

LHCII protein was extracted from spinach leaves and biochemically purified using the detergent α-DDM, as previously described.^30^ Denaturing and native gel electrophoresis indicate a high protein purity and the natural trimeric state (see **Figure S1** in the Supporting Information). We chose the small organic chromophore Texas Red (TR)^32^ as an ideal candidate for an energy donor to LHCII in membranes because of its complementary spectral properties and its amenability to assembly into lipid bilayers, using a form which is tethered to a DHPE lipid headgroup (TR-DHPE, as purchased).^33^ Ensemble absorption and fluorescence spectroscopy was performed on isolated LHCII and TR to quantify the relative optical properties in their isolated state, as a baseline for comparison to the connected system. Our LHCII sample had the expected absorption spectrum with peaks representing chlorophyll (Chl) and carotenoids between 400-500 nm and the Qy bands of Chl *b* and Chl *a* at 650 and 675nm, respectively (**Figure 1A**, *green solid*).^30^ A single fluorescence emission peak at ∼681.5 nm represents emission from lowest energy Chl *a* indicating the highly-connected chlorophyll network expected within trimeric LHCII (**Figure 1A**, *green dotted*).^30^ The absorption of Texas Red is quite broad with a maximum at 591 nm (**Figure 1A**, *solid red*), fitting well into the ‘green gap’ of LHCII. The TR fluorescence emission peak at 610 nm has extensive overlap with the LHCII Chl *b* absorption (**Figure 1A**, *dotted red*), thus the excited state of TR is energetically close to chromophores within the intended acceptor LHCII. Therefore, high efficiency donor-to-acceptor FRET would be feasible if they can be arranged into close spatial proximity, as suggested in the schematic in **Figure 1B**. The chemical structures of TR-DHPE and Chl *a* are shown in **Figure 1C** and **Figure 1D** for comparison.

**Figure 1.**
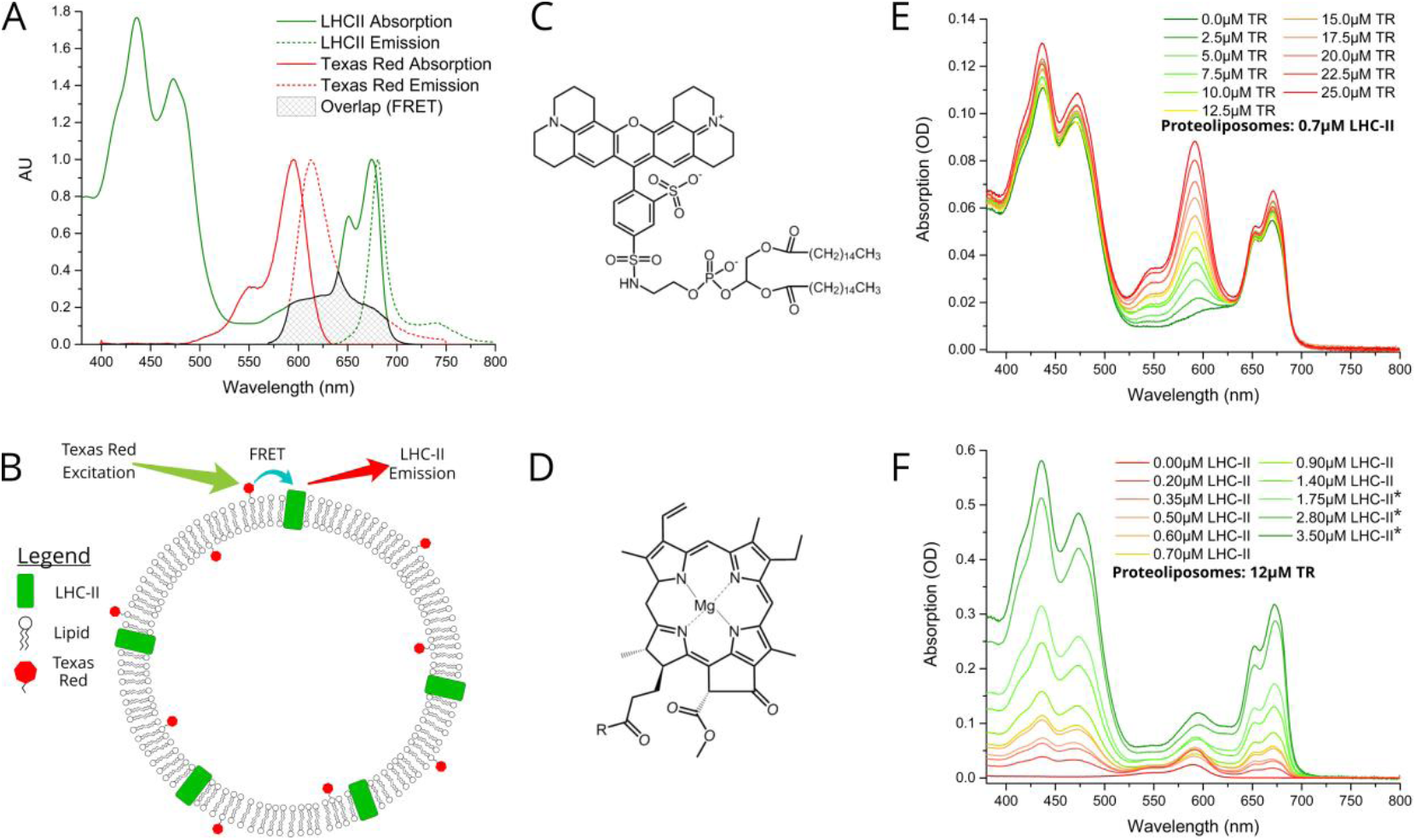
**(A)** Ensemble solution-based spectroscopy performed on dilute control samples of the isolated LHCII or TR-DHPE in a detergent suspension: normalised absorption (*solid line*) and fluorescence emission (*dotted line*) spectra. These should be thought of as the starting material and compared with the assembled proteoliposomes. LHCII was excited at 473 nm, selective excitation of Chl *b*, so that Chl *b* to Chl *a* energy transfer is monitored. TR was excited at 540 nm. *Green*: 0.4 nM LHCII trimers in a buffer of 0.03% w/v α-DDM, 10mM HEPES, pH7.5 (absorbance ∼0.1 at 675 nm). *Red*: solution of liposomes comprised of 2.1 µM TR-DHPE and 1.0 mM plant thylakoid lipids in a buffer 10mM HEPES, pH 7.5 (absorbance ∼0.1 at 561 nm). The amplitude-weighted fluorescence lifetime of these samples were 3.9 ns and 4.4 ns, respectively (fluorescence decay curves shown in **Figure S5** in the Supporting Information). **(B)** Schematic concept of proteoliposomes with energy transfer direction and wavelengths indicated (not to scale). Chemical structures of **(C)** TR-DHPE (Texas Red 1,2-dihexadecanoyl-*sn*-glycero-3-phosphoethanolamine), and **(D)** chlorophyll *a*, where R = C_20_H_40_O (phytol tail). Ensemble absorption spectra of **(E)** proteoliposome Series 1 and **(F)** proteoliposome Series 2. The *green* to *red* colour scheme represents an increasing TR-to-LHCII ratio. Proteoliposome samples were prepared and then diluted in buffer (20mM HEPES, 40mM NaCl, pH 7.5) before spectroscopy. Raw data is shown for 1/15x diluted samples, with a 3x correction factor applied to 1/45x diluted samples (those denoted by *). Proteoliposomes Series 1 had a measured TR-DHPE range from 1.7 to 21.9µM, with LHCII at 0.52-0.61µM, and 1.0mM total lipids (this equates to 2.1% TR-DHPE relative to total lipid mass, 0-35% LHCII relative to total mass). Proteoliposomes Series 2 had a measured LHCII range from 0.35 to 2.47µM, with TR-DHPE at 8.9-14.9µM, and 1.0 mM total lipids (this equates to 0-4.4% TR-DHPE relative to total lipid mass, 9.7% LHCII relative to total mass).

We designed a procedure to co-reconstitute LHCII and TR-DHPE into membrane vesicles in a modular fashion, i.e., for easy selection of the component concentrations. The protocol is similar to established proteoliposome formation procedures: initially both LHCII and lipids are fully solubilised in α-DDM detergent, followed by gradual detergent removal using porous absorptive beads (Biobeads),^26, 28^ causing the thermodynamically-driven self-assembly of membranes (see **Materials and Methods** in the Supporting Information). Analytical ultracentrifugation of proteoliposome versus control samples on Ficoll density gradients suggested that the vesicles were a consistent and homogenous single population of diameter ∼50-100nm, with no evidence of any aggregated protein (see **Figure S2** in the Supporting Information). Some general observations can be made for all proteoliposome samples as compared to the isolated LHCII and TR-DHPE: (i) the TR absorption and fluorescence peaks appear identical (for all samples), (ii) the LHCII absorption peaks have minimal shifts (<1-3 nm), and (iii) the single LHCII fluorescence peak is very similar (<1 nm shift, see **Figure S3** in the Supporting Information). This indicates that both TR-DHPE and LHCII are structurally and functionally intact within proteoliposomes. Finally, confocal fluorescence microscopy confirmed that LHCII and TR co-localised in 80-99% of the proteoliposomes in representative samples (quantified in detail below).

We wished to test whether a simple concept based on FRET could be applied to channel energy from Texas Red to LHCII in a controllable manner. From Förster theory one expects energy transfer efficiency (ETE) to be dependent on donor-acceptor distance *R*, specifically ETE∝*1/R*^*6*^ assuming incoherent energy transfer between weakly-coupled chromophores,^34^ as may be expected in our system.^35^ Thus, one should be able to modulate energy transfer with the expectation, firstly, of increasing ETE with increasing acceptor concentration and, secondly, of increasing overall magnitude of transfer with increasing donor concentration. 22 samples were prepared in two series: aiming for either a constant acceptor content (Series 1) or a constant donor content (Series 2). The LHCII and TR-DHPE composition achieved in each proteoliposome sample was quantified via absorption spectroscopy and deconvolution analysis (see **Figure S4** in the Supporting Information). Incorporation yields of both LHCII and TR were consistently high, between 75-85% and 80-90%, respectively (% of the starting material successfully incorporated). Series 1 samples had very similar LHCII concentrations of 0.52-0.61 µM with a TR range from 1.7 to 21.9 µM (**Figure 1E**). Series 2 samples had relatively similar TR concentrations of 8.9-14.9 µM with a LHCII range from 0.35 to 2.47µM (**Figure 1F**). Generation of this proteoliposome composition range demonstrates the modularity of our system in terms of control over the incorporation of LHCII and TR-DHPE.

A well-defined system would be one where additional energy can be channelled to LHCII from extrinsic donors in a controllable manner. To estimate FRET efficiency the fluorescence of the donor chromophore in the combined system (i.e. donor + acceptor) was compared to the donor-only system (i.e. isolated donors). Donor-to-acceptor energy transfer causes a quenching of the donor fluorescence, manifested as decreased emission intensity concomitant with a faster decay of excited state, due to a portion of the donor excitons following this alternative non-radiative transfer pathway.^20^ For our samples, the relative TR fluorescence intensity (TR emission per mole TR) was quantified in steady-state fluorescence emission spectra after selective excitation of TR at 540nm. Consideration of proteoliposome Series 2 (LHCII range at constant TR) allowed a quantitative analysis of how TR quenching depends upon LHCII concentration. A clear qualitative trend of decreasing TR fluorescence intensity with increasing LHCII concentration was observed, as energy is increasingly transferred from TR to LHCII by FRET (**Figure 2A**). Relative TR fluorescence as low as 2% of the isolated TR-DHPE control is found in proteoliposomes where LHCII concentration is at the maximum tested (2.84 μM). Time-resolved fluorescence measurements were also performed as a secondary technique to independently quantify TR quenching, with selective excitation of TR using a picosecond-pulsed 540 nm laser. Fluorescence decay curves of proteoliposome Series 2 revealed a faster decay of TR fluorescence as LHCII concentration was increased (**Figure 2C**), representing quenching in correlation with the steady-state data. Fitting of the curves to an exponential decay function shows that the TR lifetime decreases from 4.4 ns in the absence of LHCII to a minimum of 0.7 ns at maximum LHCII concentration (2.84 μM). As a further test, we took measurements before and after complete re-solubilisation of the membranes with excess α-DDM detergent. Re-solubilisation resulted in regeneration of highly fluorescent TR, confirming that its quenching is dependent on the membrane architecture for allowing interaction with LHCII (see **Figure S6** in the Supporting Information).

**Figure 2.**
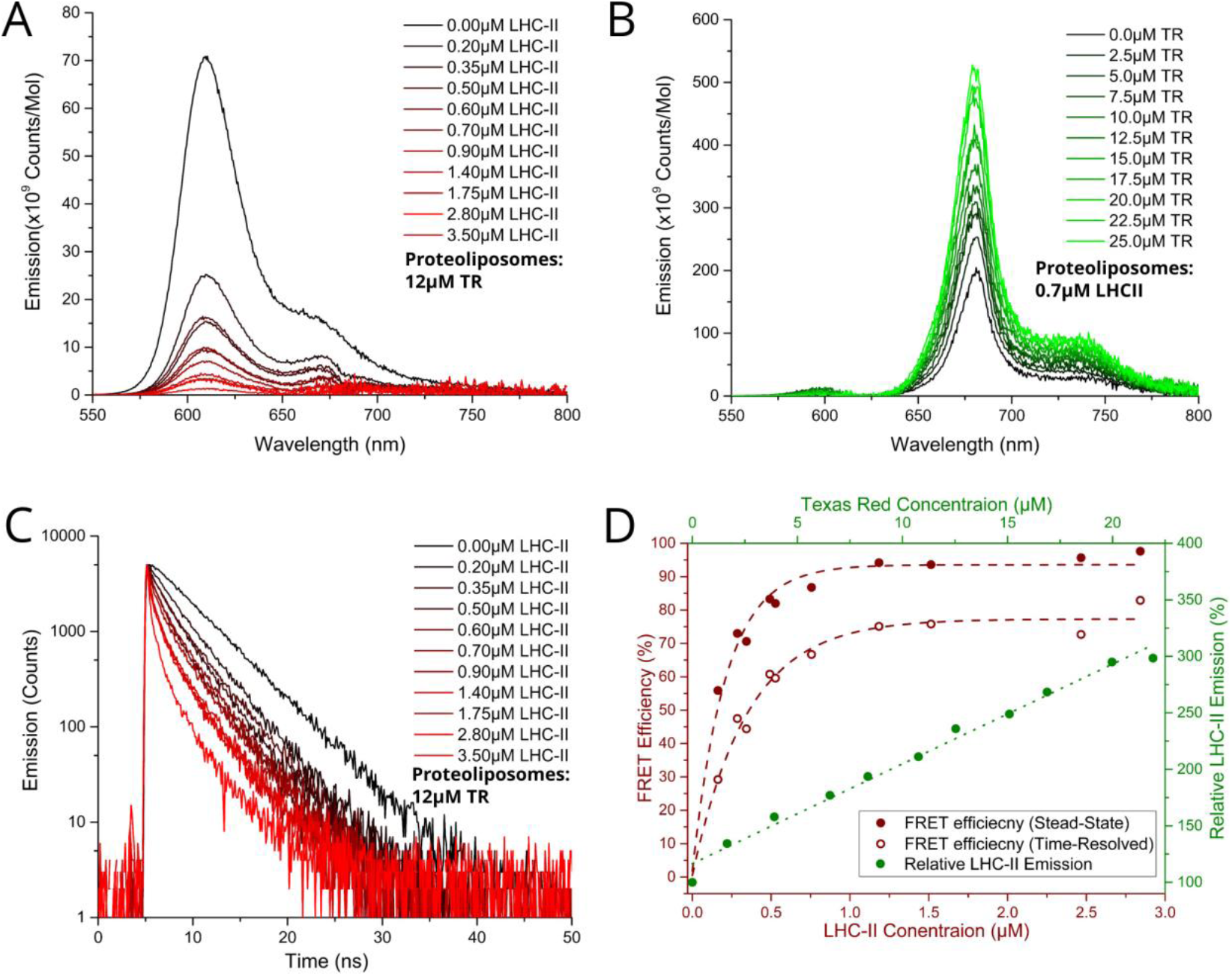
**(A)** Steady-state fluorescence emission spectra of sample Series 2 with selective excitation of TR at 540 nm. For clarity, spectra were been de-convoluted to subtract the LHCII peak (see **Figure S7** in the Supporting Information). **(B)** Steady-state fluorescence emission of sample Series 1 with selective excitation TR at 540 nm. For clarity, spectra were de-convoluted to subtract the TR peak. **(C)** Time-resolved fluorescence measurements of sample Series 2 with TR excitation by a 540 nm pulsed laser. The full dataset of fluorescence intensities and lifetimes are given in **Table S1** in the Supporting Information. **(D)** *Red labelling*: Energy Transfer Efficiency versus LHCII concentration, calculated from proteoliposome Series 2 (bottom and left axis). Energy transfer efficiency was calculated from both steady-state (ETE_SS_) and time-resolved (ETE_TR_) data by comparison of samples with and without acceptors (LHCII) using the conventional relationship for FRET between donor *D* and acceptor *A*:

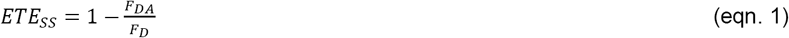

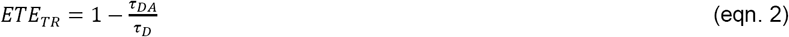

where F_DA_ and F_D_ are the relative fluorescence intensity of TR in the presence or absence of LHCII, respectively, and*τ*_DA_ and *τ*_D_ are the mean fluorescence lifetime of TR the presence or absence of LHCII, respectively. **(D)** *Green labelling*: Relative LHCII emission versus TR concentration, calculated from proteoliposome Series 1 (top and right axis). Note, the known phenomenon of self-quenching of LHCII fluorescence, which increases with LHCII content in proteoliposomes,^24, 25, 28^ was quantified in control samples omitting TR-DHPE (see **Figure S8** in the Supporting Information). This limits the absolute intensity of LHCII emission as compared to detergent-solubilized LHCII. Here, 100% is defined as the baseline intensity of LHCII emission observed for proteoliposomes containing 0.7μM LHCII and 1 mM total lipid because this is the average LHCII content in proteoliposome Series 1. The possibility of self-quenching of TR fluorescence was also considered and found to be minimal at 12µM TR (see **Figure S9** in the Supporting Information).

Energy transfer from TR to LHCII was also apparent from observed increases in the relative LHCII fluorescence intensity (LHCII emission per mole LHCII) after selective excitation of TR at 540nm. LHCII had a quantifiable low level of fluorescence without TR, but this increased significantly with increasing TR-DHPE concentration (**Figure 2B**). In proteoliposomes containing the highest TR-DHPE content investigated (21.9 μM) the LHCII emission reached a maximum of 3.14x enhancement relative to proteoliposomes without TR-DHPE.

Graphical analysis of fluorescence intensity and fluorescence lifetime data allowed the observed trends to be quantified. Energy transfer efficiency was calculated from both steady-state (ETESS) and time-resolved (ETETR) data by comparison of samples with and without acceptors (LHCII). **Figure 2D** (*red datapoints*) shows that efficiency increased non-linearly with LHCII concentration and is fitted to an exponential growth function, as efficiency is expected to saturate at high acceptor concentration. Steady-state and time-resolved data show good agreement of the trend and estimate a maximal ETE of ∼94% and ∼77%, respectively. We expect that the exact maximal ETE is between these two values (different spectroscopic techniques often differ in absolute efficiency values, providing a definitive calculation of ETE will be studied in future work). This trend of ETE is attributed simply to the reduction in average distance between TR and LHCII as LHCII concentration is increased. The relative LHCII emission was found to increase in proportion to the concentration of TR-DHPE, for all concentrations investigated, see **Figure 2D** *green datapoints*. This linear relationship is consistent with a constant TR-to-LHCII energy transfer efficiency irrespective of TR-DHPE concentration. Note that LHCII self-quenching and TR self-quenching was quantified in control samples and taken into account (see legend in **Figure 2**). In conclusion, the proteoliposome system has well-defined donor-to-acceptor energy transfer and TR provides a controllable enhancement of LHCII emission demonstrating the benefit of increased spectral coverage in the ‘green gap’.

Assessment of proteoliposome functionality when associated with planar surfaces is important for any future applications requiring thin films (e.g. in photo-active devices). Furthermore, microscopy is complementary to ensemble spectroscopy, allowing the identification and analysis of individual proteoliposomes. Fluorescence Lifetime Imaging Microscopy (FLIM) was performed on two representative samples deposited onto glass coverslips in order to investigate population differences in samples with different energy transfer efficiencies: (i) “low-LHCII” proteoliposomes (low ETE) and (ii) “high-LHCII” proteoliposomes (high ETE) with equal concentrations of TR (1.2 µM LHCII + 12 µM TR-DHPE and 2.8 µM LHCII + 12 µM TR-DHPE, respectively). TR and LHCII were selectively excited with 561 nm and 485 nm pulsed interleaved lasers, respectively, and observed in dedicated emission channels simultaneously (see **Materials and Methods** in the Supporting Information). This allowed visualisation of fluorescence intensity and fluorescent lifetimes for both TR and LHCII within individual proteoliposomes. Hundreds of well-resolved particles in each 25 × 25 µm field of view were observed with apparent correlation in both TR and LHCII channel intensity images (**Figure 3A**). Quantitative analysis of these sample is non-trivial as they are weakly emitting, subject to photo-damage, self-quenching and inter-molecule energy transfer is taking place. Therefore, the effects of spectral overlap, photo-bleaching, detector noise and potential contaminants were quantified and subtracted from all particles (see **Materials and Methods** in the Supporting Information). A population analysis of single proteoliposomes was performed by manually selecting particles within images, tabulating the data of fluorescence intensity and lifetimes, and performing statistical analysis. These manual analyses allow the LHCII and TR to be independently identified in individual proteoliposomes revealing the relationship between the two fluorescent components. The fraction of proteoliposomes which contain both TR and LHCII was estimated. At least 83% and 80% of proteoliposomes had significant signal from both TR and LHCII emission channels, for low-LHCII and high-LHCII samples, respectively (n= 200, with 99% confidence relative to background fluorescence, see **Table S2** in the Supporting Information). This may underestimate the presence of LHCII, as smaller vesicles or those with a lesser quantity of LHCII may not be detected.

**Figure 3.**
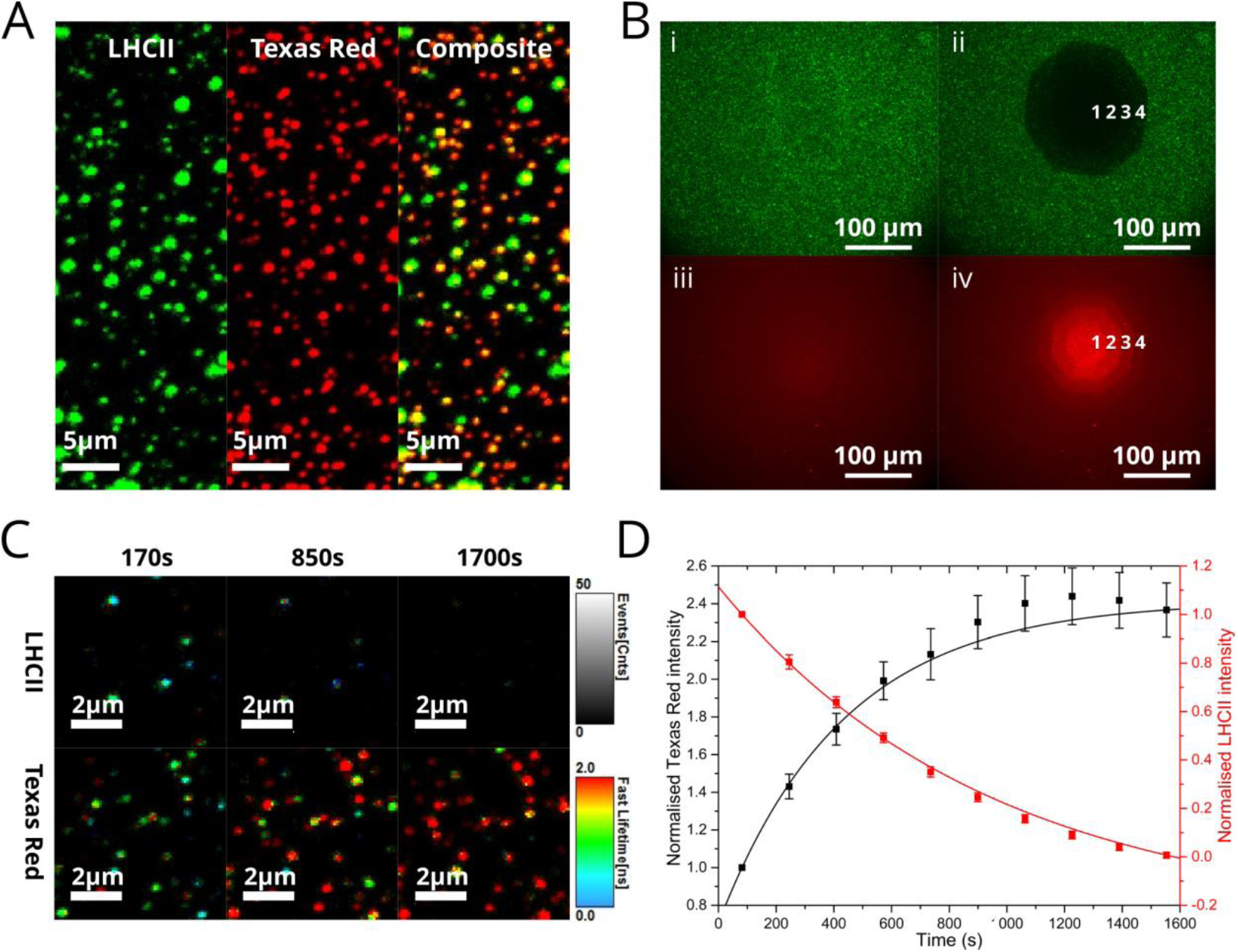
Fluorescence microscopy and photo-bleaching data from proteoliposomes deposited on glass at either low or high surface density, prepared by incubating clean hydrophilic glass coverslips with proteoliposome samples diluted with buffer as required. All microscopy was performed in a buffer of 20mM HEPES, 40mM NaCl (pH 7.5). See **Materials and Methods** in the Supporting Information for full details of the image acquisition and analysis for all parts. **(A)** FLIM images of a representative field of view of proteoliposomes at low surface density, panels showing fluorescence intensity recorded in either *LHCII channel* or *TR channel*, and a *Composite* overlay of both images. Strong co-localisation of signals from LHCII and TR leads to a yellow colour. Images shown here are from a proteoliposome sample containing 2.8μM LHCII, 12μM TR, 1 mM total lipid; a gallery of images from both samples assessed by FLIM is shown in **Figure S10** in the Supporting Information. **(B)** Epifluorescence microscopy images of a representative proteoliposome sample (3.5μM LHCII, 12 µM TR, 1 mM total lipid) deposited at high surface density. *LHCII channel* and *TR channel* were recorded sequentially using appropriate filter cubes. Bleaching was performed by exposing the sample to a high intensity of LHCII excitation through an aperture that is consecutively opened in 4 steps over 90 s. **(C)** A time-lapse series of FLIM images of proteoliposomes where LHCII photo-bleaching occurs across the whole image, with minimal TR bleaching in comparison, displaying both fluorescence intensity (*brightness*) and “fastFLIM lifetime” (*colour scale*). The “fastFLIM lifetime” is the mean time of photon arrival, a good approximation of the expected fitted lifetime, which allowed high throughput multi-image analysis and circumvents the challenge of fitting decay curves where photon counts are low. **(D)** Graph showing the total fluorescence intensity of the *LHCII* and *TR channels* versus time, for the sequence from (C).

Photo-bleaching experiments were performed using epifluorescence microscopy to provide a visual representation of energy transfer at the microscale, allowing us to assess whether optical functionality is retained when proteoliposomes are at high density on surfaces. A confined region of the sample was exposed to a high intensity of light in order to photo-bleach LHCII. Selective photo-bleaching of LHCII was performed using appropriate filters and the effects observed in subsequent fluorescence images. A very low LHCII fluorescence was observed in the bleach region (**Figure 3B**, *(i)* vs *(ii)*), whereas, the TR fluorescence in this region showed an apparent increase (**Figure 3B**, *(iii)* vs *(iv)*). We could vary the degree of LHCII photo-bleaching and demonstrate the concomitant TR de-quenching by iteratively opening the aperture in stages (**Figure 3B**, *numbers 1-4*). This “de-quenching” of TR fluorescence confirms that photo-damaged LHCII is unable to act as an energy acceptor from Texas Red. FLIM was used to analyse the photo-bleaching of proteoliposomes at lower surface densities and higher resolution. Bleaching occurs across the entire image region during continuous FLIM acquisition, but LHCII bleaches much faster than TR, allowing us to distinguish these effects. A decrease in LHCII fluorescence intensity with a concomitant increase in TR intensity is apparent in images of proteoliposomes in **Figure 3C** and clear when plotting a time-course of the fluorescence intensity, **Figure 3D.** This FLIM data is a dynamic representation of the same photo-bleaching effects observed in epifluorescence data in **Figure 3B**. The same trend is apparent in time-resolved data, where the estimated TR lifetimes increase over time. **Supplementary Video 1** (in the Supporting Information) shows this particularly clearly. By monitoring individual proteoliposomes across multiple images the fraction of proteoliposomes in which FRET occurs can be determined by the proxy of the TR de-quenching effect. Approximately 90% and 95% of proteoliposomes displayed an increase in TR lifetime (≥225% between the start and end of the FLIM acquisition), for low-LHCII and high-LHCII samples, respectively. The mean TR lifetime increased from 0.98 ±0.54 ns to 2.75 ±0.71 ns for ‘high-LHCII’ and from 1.15 ±0.70 ns to 2.77 ±0.76 ns for ‘low-LHCII’ (n=100 for both). In summary, the photo-bleaching data from both epifluorescence and FLIM conclusively shows that the vast majority of proteoliposomes contain both LHCII and TR and that energy transfer between them is retained when these membranes are deposited onto a surface.

In order to calculate the fluorescence lifetimes on a per-proteoliposome basis, manual analysis was performed by fitting fluorescence decay curves extracted from a region-of-interest defined to represent individual proteoliposomes. Only well-resolved proteoliposomes with sufficient signal to produce a good fit were selected (criteria of counts >500 and fit chi^2^ <1.2). Fluorescence decay curves from five representative proteoliposomes selected from a field in **Figure 4A** are displayed in **Figure 4B**, alongside calculated mean lifetimes of both TR and LHCII components. Frequency distributions of the fitted TR lifetime for different proteoliposome samples (**Figure 4C**) indicate that TR fluorescence is quenched by LHCII. The mean TR fluorescence lifetime decreases from 3.26 ± 0.46 ns in control TR liposomes (without LHCII) to 1.79 ± 0.64 ns and 1.46 ± 0.53 ns for “low-LHCII” and “high-LHCII” proteoliposomes, when integrating over 500 frames of data. This long acquisition provides high enough signal for good fitting, but (again) causes photo-bleaching of LHCII resulting in a distortion of the TR lifetime towards larger values. So, the energy transfer efficiency (ETE) calculated after correction for this bleaching was plotted as a frequency distribution (**Figure 4D**). We find a mean ETE ±S.D. of 71 ±10% and 77 ±8% for “low-LHCII” and “high-LHCII” proteoliposomes, respectively. This shows that within a population, the ETE varies between individual proteoliposomes. It is interesting to note that a significant fraction of vesicles has an ETE >90% and we speculate that it may be possible to biochemically purify a sub-population of vesicles enriched for high ETE.^23, 36^ In summary, the FLIM population distributions confirm that FRET occurs with increasing ETE with increasing LHCII concentration, in agreement with our ensemble spectroscopy data. Importantly, our single-particle FLIM analyses prove that our proteoliposomes are have quantitatively similar FRET on surfaces as in solution in bulk. Our study is one of the few where single-molecule spectroscopy was utilised to quantify distributions of fluorescence lifetime in membrane reconstituted light-harvesting systems^25^ beyond traditional ensemble spectroscopy. Observing the heterogeneity in this way could reveal low-frequency events and also guide the optimization of the system if one wishes to maximize any aspect, e.g. transfer efficiency.

**Figure 4.**
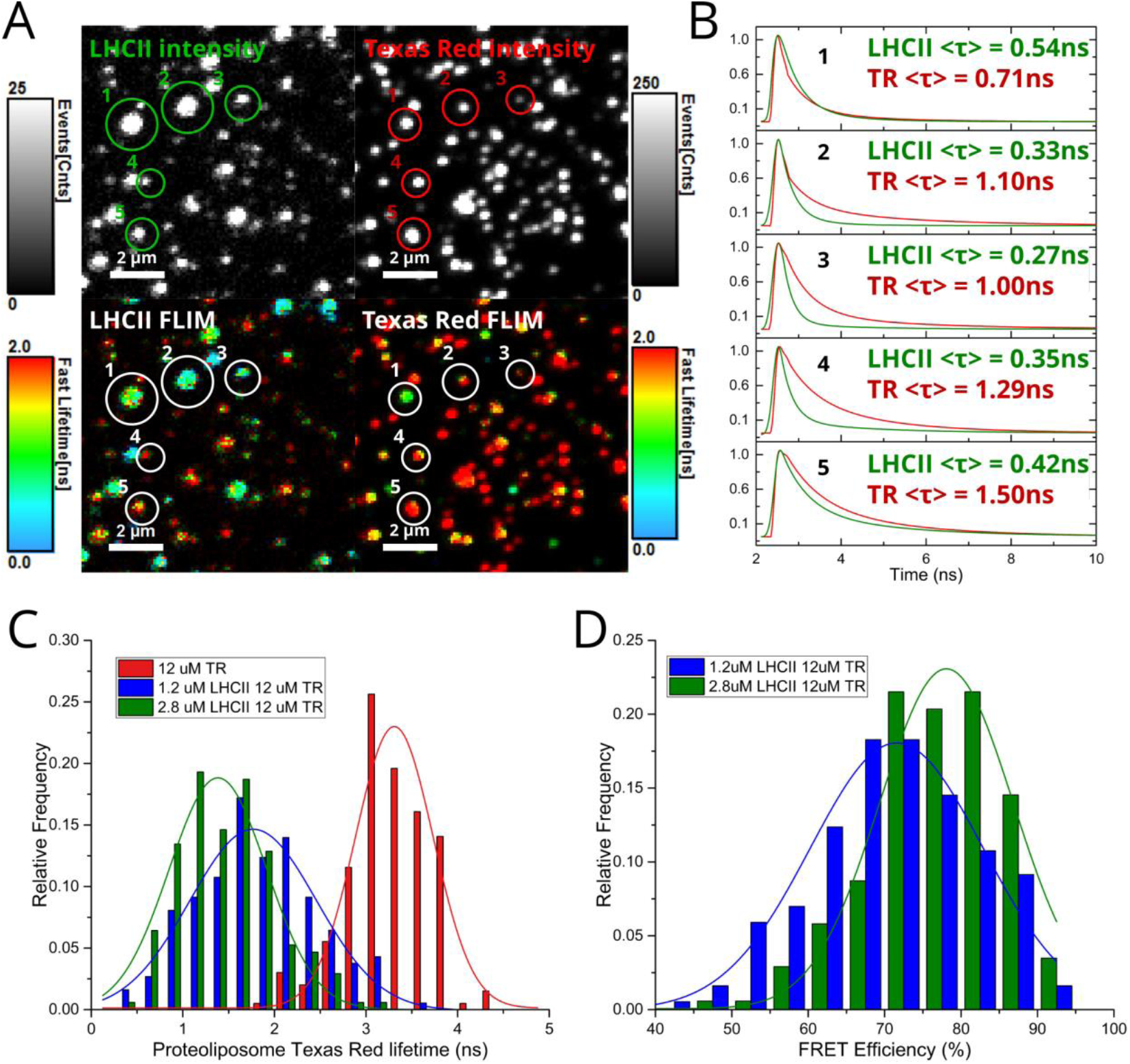
FLIM data from proteoliposomes deposited at a low surface density on glass coverslips. See **Materials and Methods** in the Supporting Information for full details of the analysis for all parts. **(A)** A representative field of proteoliposomes imaged at high magnification, comparing fluorescence intensity (*top*) versus fluorescence lifetime (*bottom*) between LHCII and TR channels, as labelled. Five representative particles have been indicated *1-5*. Images from the proteoliposome sample containing 2.8μM LHCII + 12μM TR, 1 mM total lipid. **(B)** Fluorescence decay curve fits from the individual proteoliposomes, as numbered in (A), annotated with calculated mean amplitude-weighted lifetimes <***τ***>, as labelled for each components. Here, the fluorescence data from the 20-30 pixels representing one proteoliposome were binned, a decay curve was generated, and re-convolution fit to a biexponential decay function was performed. **(C)** Frequency distribution histogram of the mean TR fluorescence lifetime, comparing the low-LHCII proteoliposome, high-LHCII proteoliposome and TR liposome samples (n = 186, 175 and 200, respectively, see **Table S2** in the Supporting Information). These TR lifetimes are overestimates due to the LHCII bleaching and subsequent de-quenching of TR due to LHCII photo-bleaching during the acquisition period. **(D)** Frequency distribution histogram of the ETE comparing the proteoliposome sample, calculated as in *Figure 2 equation (2)* using the data from (C). A correction factor determined from control samples is applied to account for de-quenching of TR (see **Table S3** in the Supporting Information).

In conclusion, using a proteoliposome-based system we have significantly enhanced the effective absorption range of LHCII via energy transfer from the spectrally complementary chromophore Texas Red. Specifically, by exploiting the fact that lipids and membrane proteins self-assemble into nano-sized architectures the distance between TR and LHCII is shortened, whereas, when not confined these nanoscale membranes (e.g. when solubilised by detergent) the energy transfer drops to almost zero. Energy transfer between TR and LHCII in proteoliposomes was highly efficient, comparable to systems utilising covalent attachment of chromophores to the protein,^11-14^ and is consistent with theoretical modelling of a system that has a random distribution of chromophores throughout a 2D lipid bilayer undergoing simple donor-to-acceptor FRET.^37^ This, together with our single-proteoliposome FLIM data, suggests that our self-assembly protocol creates a well-mixed environment on the nanoscale, addressing the challenge of inconsistent distribution of membrane proteins between proteoliposomes that was previously reported with other similar methods,^23^ with the potential that further purification could isolate sub-populations with desirable properties.^36^ We demonstrated the modularity of these membranes by controlling the concentration of both LHCII and TR-DHPE as desired across an order of magnitude. Future studies could incorporate a suite of different organic chromophores and/or quantum dots into lipid bilayers to interact with LH proteins to enhance its spectral range even further. Alternatively these membranes have potential to improve our understanding of the photophysical properties of LH proteins, for example, the transient interactions between diffusing chromophores and LH proteins could be assessed using ultrafast spectroscopy.^13, 38^ Our group previously demonstrated diblock copolymer assemblies can also be used as a matrix to arrange organic chromophores for high efficiency FRET^20-22^ and may be more robust than lipid bilayers, however, they are not a natural environment for membrane proteins. We speculate that a polymer-lipid hybrid^39, 40^ system could be tuned to have the advantages of a more robust polymeric system and protein compatibility of lipids.

Our LHCII/TR proteoliposomes were found to retain their energy transfer efficiency when deposited onto solid supports, providing proof-of-principle that proteoliposomes can be used for surface-based applications. Previous studies have shown that LH proteins attached to glass surfaces in nanoscale array patterns provide directional transfer of energy^41, 42^ and LH proteins on nanostructured gold can modulate surface plasmon resonances.^43^ Expanding on these ideas to make use of our LHCII/TR proteoliposome methodology, future studies could pattern membranes which include any LH proteins and additional chromophores as desired,^33^ for the generation of surface-supported organized nanoscale membranes which have enhanced light-harvesting capability. If our proteoliposomes form well-connected thin films on surfaces (preliminary characterization by atomic force microscopy is underway) they could have future applications as photo-active bio-hybrid materials, for example as coatings to enhance the current generated by photoelectrochemical devices.^44, 45^

## Supporting information

Video 1 - LHCII

## Acknowledgements

A.M.H. was supported by an Engineering and Physical Sciences Research Council (EPSRC, UK) studentship, award number 1807029, and an EPSRC programme grant, EP/J017566/1 ‘‘CAPITALs’’. S.A.M. was supported by a Biotechnology and Biological Sciences Research Council (BBSRC, UK) studentship, award number 1940236. S.D.A.C. was also supported by EPSRC ‘‘CAPITALs’’ award EP/J017566/1. L.J.C.J. is grateful to the BBSRC for funding (BB/P005454/1). P.G.A. was supported by a Future Leader Fellowship from the BBSRC, award number BB/M013723/1, and a University Academic Fellowship (University of Leeds). The PicoQuant FLIM instrument was acquired at Leeds with funding from a BBSRC award number BB/R000174/1, and the Quantamaster fluorescence spectrometer was funded by BBSRC award number BB/R000271/1.

P.G.A. thanks Eliot Dawson (Leeds) for the preliminary work he performed on this project.

## References

1. Gust, D.; Moore, T. A.; Moore, A. L., Solar Fuels via Artificial Photosynthesis, Accounts of Chemical Research 2009, 42, (12), 1890–1898.

2. Li, X. M.; Qiao, S. P.; Zhao, L. L.; Liu, S. D.; Li, F.; Yang, F. H.; Luo, Q.; Hou, C. X.; Xu, J. Y.; Liu, J. Q., Template-Free Construction of Highly Ordered Monolayered Fluorescent Protein Nanosheets: A Bioinspired Artificial Light-Harvesting System, ACS Nano 2019, 13, (2), 1861–1869.

3. Berhanu, S.; Ueda, T.; Kuruma, Y., Artificial photosynthetic cell producing energy for protein synthesis, Nature Communications 2019, 10, 1325.

4. Liu, C.; Gallagher, J. J.; Sakimoto, K. K.; Nichols, E. M.; Chang, C. J.; Chang, M. C. Y.; Yang, P. D., Nanowire-Bacteria Hybrids for Unassisted Solar Carbon Dioxide Fixation to Value-Added Chemicals, Nano Letters 2015, 15, (5), 3634–3639.

5. Qiu, J.; Zeng, G. T.; Ha, M. A.; Ge, M. Y.; Lin, Y. J.; Hettick, M.; Hou, B. Y.; Alexandrova, A. N.; Javey, A.; Cronin, S. B., Artificial Photosynthesis on TiO2-Passivated InP Nanopillars, Nano Letters 2015, 15, (9), 6177–6181.

6. Sun, H. C.; Zhang, X. Y.; Miao, L.; Zhao, L. L.; Luo, Q.; Xu, J. Y.; Liu, J. Q., Micelle-Induced Self-Assembling Protein Nanowires: Versatile Supramolecular Scaffolds for Designing the Light-Harvesting System, ACS Nano 2016, 10, (1), 421–428.

7. Blankenship, R. E.; Tiede, D. M.; Barber, J.; Brudvig, G. W.; Fleming, G.; Ghirardi, M.; Gunner, M. R.; Junge, W.; Kramer, D. M.; Melis, A.; Moore, T. A.; Moser, C. C.; Nocera, G.; Nozik, A. J.; Ort, D. R.; Parson, W. W.; Prince, R. C.; Sayre, R. T., Comparing Photosynthetic and Photovoltaic Efficiencies and Recognizing the Potential for Improvement, Science 2011, 332, (6031), 805–809.

8. Gao, J. L.; Wang, H.; Yuan, Q. P.; Feng, Y., Structure and Function of the Photosystem Supercomplexes, Frontiers in Plant Science 2018, 9, 357.

9. Mirkovic, T.; Ostroumov, E. E.; Anna, J. M.; van Grondelle, R.; Govindjee; Scholes, G. D., Light Absorption and Energy Transfer in the Antenna Complexes of Photosynthetic Organisms, Chemical Reviews 2017, 117, (2), 249–293.

10. Standfuss, J.; van Scheltinga, A. C. T.; Lamborghini, M.; Kuhlbrandt, W., Mechanisms of photoprotection and nonphotochemical quenching in pea light-harvesting complex at 2.5 Å resolution, EMBO Journal 2005, 24, (5), 919–928.

11. Gundlach, K.; Werwie, M.; Wiegand, S.; Paulsen, H., Filling the “green gap” of the major light-harvesting chlorophyll a/b complex by covalent attachment of Rhodamine Red, Biochimica et Biophysica Acta, Bioenergetics 2009, 1787, (12), 1499–504.

12. Harris, M. A.; Jiang, J.; Niedzwiedzki, D. M.; Jiao, J.; Taniguchi, M.; Kirmaier, C.; Loach, P. A.; Bocian, D. F.; Lindsey, J. S.; Holten, D.; Parkes-Loach, P. S., Versatile design of biohybrid light-harvesting architectures to tune location, density, and spectral coverage of attached synthetic chromophores for enhanced energy capture, Photosynthesis Research 2014, 121, (1), 35–48.

13. Yoneda, Y.; Noji, T.; Katayama, T.; Mizutani, N.; Komori, D.; Nango, M.; Miyasaka, H.; Itoh, S.; Nagasawa, Y.; Dewa, T., Extension of Light-Harvesting Ability of Photosynthetic Light-Harvesting Complex 2 (LH2) through Ultrafast Energy Transfer from Covalently Attached Artificial Chromophores, Journal of the American Chemical Society 2015, 137, (40), 13121–9.

14. Springer, J. W.; Parkes-Loach, P. S.; Reddy, K. R.; Krayer, M.; Jiao, J.; Lee, G. M.; Niedzwiedzki, D. M.; Harris, M. A.; Kirmaier, C.; Bocian, D. F.; Lindsey, J. S.; Holten, D.; Loach, P. A., Biohybrid Photosynthetic Antenna Complexes for Enhanced Light-Harvesting, Journal of the American Chemical Society 2012, 134, (10), 4589–4599.

15. Schmitt, F. J.; Maksimov, E. G.; Hätti, P.; Weißenborn, J.; Jeyasangar, V.; Razjivin, A. P.; Paschenko, V. Z.; Friedrich, T.; Renger, G., Coupling of different isolated photosynthetic light harvesting complexes and CdSe/ZnS nanocrystals via Förster resonance energy transfer, Biochimica et Biophysica Acta, Bioenergetics 2012, 1817, (8), 1461–1470.

16. Werwie, M.; Xu, X.; Haase, M.; Basche, T.; Paulsen, H., Bio Serves Nano: Biological Light-Harvesting Complex as Energy Donor for Semiconductor Quantum Dots, Langmuir 2012, 28, (13), 5810–5818.

17. Sahin, T.; Harris, M. A.; Vairaprakash, P.; Niedzwiedzki, D. M.; Subramanian, V.; Shreve, A. P.; Bocian, D. F.; Holten, D.; Lindsey, J. S., Self-Assembled Light-Harvesting System from Chromophores in Lipid Vesicles, The Journal of Physical Chemistry. B 2015, 119, (32), 10231–43.

18. De Leo, V.; Catucci, L.; Falqui, A.; Marotta, R.; Striccoli, M.; Agostiano, A.; Comparelli, R.; Milano, F., Hybrid Assemblies of Fluorescent Nanocrystals and Membrane Proteins in Liposomes, Langmuir 2014, 30, (6), 1599–1608.

19. Lukashev, E. P.; Knox, P. P.; Gorokhov, V. V.; Grishanova, N. P.; Seifullina, N. K.; Krikunova, M.; Lokstein, H.; Paschenko, V. Z., Purple-bacterial photosynthetic reaction centers and quantum-dot hybrid-assemblies in lecithin liposomes and thin films, Journal of Photochemistry and Photobiology B: Biology 2016, 164, (Supplement C), 73–82.

20. Adams, P. G.; Collins, A. M.; Sahin, T.; Subramanian, V.; Urban, V. S.; Vairaprakash, P.; Tian, Y.; Evans, D. G.; Shreve, A. P.; Montano, G. A., Diblock copolymer micelles and supported films with noncovalently incorporated chromophores: a modular platform for efficient energy transfer, Nano Letters 2015, 15, (4), 2422–8.

21. Orf, G. S.; Collins, A. M.; Niedzwiedzki, D. M.; Tank, M.; Thiel, V.; Kell, A.; Bryant, D. A.; Montano, G. A.; Blankenship, R. E., Polymer-Chlorosome Nanocomposites Consisting of Non-Native Combinations of Self-Assembling Bacteriochlorophylls, Langmuir 2017, 33, (25), 6427–6438.

22. Collins, A. M.; Timlin, J. A.; Anthony, S. M.; Montano, G. A., Amphiphilic block copolymers as flexible membrane materials generating structural and functional mimics of green bacterial antenna complexes, Nanoscale 2016, 8, (32), 15056–15063.

23. Tutkus, M.; Akhtar, P.; Chmeliov, J.; Gorfol, F.; Trinkunas, G.; Lambrev, P. H.; Valkunas, L., Fluorescence Microscopy of Single Liposomes with Incorporated Pigment-Proteins, Langmuir 2018, 34, (47), 14410–14418.

24. Moya, I.; Silvestri, M.; Vallon, O.; Cinque, G.; Bassi, R., Time-Resolved Fluorescence Analysis of the Photosystem II Antenna Proteins in Detergent Micelles and Liposomes, Biochemistry 2001, 40, 12552–12561.

25. Natali, A.; Gruber, J. M.; Dietzel, L.; Stuart, M. C.; van Grondelle, R.; Croce, R., Light-harvesting Complexes (LHCs) Cluster Spontaneously in Membrane Environment Leading to Shortening of Their Excited State Lifetimes, The Journal of Biological Chemistry 2016, 291, (32), 16730–9.

26. Seiwert, D.; Witt, H.; Ritz, S.; Janshoff, A.; Paulsen, H., The Nonbilayer Lipid MGDG and the Major Light-Harvesting Complex (LHCII) Promote Membrane Stacking in Supported Lipid Bilayers, Biochemistry 2018, 57, (15), 2278–2288.

27. Zhou, F.; Liu, S.; Hu, Z.; Kuang, T.; Paulsen, H.; Yang, C., Effect of monogalactosyldiacylglycerol on the interaction between photosystem II core complex and its antenna complexes in liposomes of thylakoid lipids, Photosynthesis Research 2009, 99, (3), 185–193.

28. Crisafi, E.; Pandit, A., Disentangling protein and lipid interactions that control a molecular switch in photosynthetic light harvesting, Biochimica et Biophysica Acta, Biomembranes 2017, 1859, (1), 40–47.

29. Pandit, A.; Shirzad-Wasei, N.; Wlodarczyk, L. M.; van Roon, H.; Boekema, E. J.; Dekker, J. P.; de Grip, W. J., Assembly of the major light-harvesting complex II in lipid nanodiscs, Biophysical Journal 2011, 101, (10), 2507–15.

30. Adams, P. G.; Vasilev, C.; Hunter, C. N.; Johnson, M. P., Correlated fluorescence quenching and topographic mapping of Light-Harvesting Complex II within surface-assembled aggregates and lipid bilayers, Biochimica et Biophysica Acta, Bioenergetics 2018, 1859, (10), 1075–1085.

31. van Oort, B.; Marechal, A.; Ruban, A. V.; Robert, B.; Pascal, A. A.; de Ruijter, N. C.; van Grondelle, R.; van Amerongen, H., Different crystal morphologies lead to slightly different conformations of light-harvesting complex II as monitored by variations of the intrinsic fluorescence lifetime, Physical Chemistry Chemical Physics 2011, 13, (27), 12614–22.

32. Titus, J. A.; Haugland, R.; Sharrow, S. O.; Segal, D. M., Texas red, a hydrophilic, red-emitting flourophore for use with flourescein in dual parameter flow microfluorometric and fluorescence microscopic studies, Journal of Immunological Methods 1982, 50, (2), 193–204.

33. Adams, P. G.; Swingle, K. L.; Paxton, W. F.; Nogan, J. J.; Stromberg, L. R.; Firestone, M. A.; Mukundan, H.; Montaño, G. A., Exploiting lipopolysaccharide-induced deformation of lipid bilayers to modify membrane composition and generate two-dimensional geometric membrane array patterns, Scientific Reports 2015, 5, 10331.

34. Förster, T., Delocalized excitation and excitation transfer, Modern Quantum Chemistry Istanbul Lectures 1965, 3, 93–137.

35. Fox, K. F.; Balevicius, V.; Chmeliov, J.; Valkunas, L.; Ruban, A. V.; Duffy, C. D. P., The carotenoid pathway: what is important for excitation quenching in plant antenna complexes?, Physical Chemistry Chemical Physics 2017, 19, (34), 22957–22968.

36. Akhtar, P.; Görföl, F.; Garab, G.; Lambrev, P. H., Dependence of chlorophyll fluorescence quenching on the lipid-to-protein ratio in reconstituted light-harvesting complex II membranes containing lipid labels, Chemical Physics 2019, 522, 242–248.

37. Subramanian, V.; Zurek, N. A.; Evans, D. G.; Shreve, A. P., Predictive modeling of broad wavelength light-harvesting performance in assemblies of multiple chromophores, Journal of Photochemistry and Photobiology A: Chemistry 2018, 367, 105–114.

38. Ogren, J. I.; Tong, A. L.; Gordon, S. C.; Chenu, A.; Lu, Y.; Blankenship, R. E.; Cao, J.; Schlau-Cohen, G. S., Impact of the lipid bilayer on energy transfer kinetics in the photosynthetic protein LH2, Chemical Science 2018, 9, (12), 3095–3104.

39. Seneviratne, R.; Khan, S.; Moscrop, E.; Rappolt, M.; Muench, S. P.; Jeuken, L. J. C.; Beales, P. A., A reconstitution method for integral membrane proteins in hybrid lipid-polymer vesicles for enhanced functional durability, Methods 2018, 147, 142–149.

40. Otrin, L.; Marusic, N.; Bednarz, C.; Vidakovic-Koch, T.; Lieberwirth, I.; Landfester, K.; Sundmacher, K., Toward Artificial Mitochondrion: Mimicking Oxidative Phosphorylation in Polymer and Hybrid Membranes, Nano Letters 2017, 17, (11), 6816–6821.

41. Vasilev, C.; Johnson, M. P.; Gonzales, E.; Wang, L.; Ruban, A. V.; Montano, G.; Cadby, A. J.; Hunter, C. N., Reversible Switching between Nonquenched and Quenched States in Nanoscale Linear Arrays of Plant Light-Harvesting Antenna Complexes, Langmuir 2014, 30, (28), 8481–8490.

42. Escalante, M.; Lenferink, A.; Zhao, Y.; Tas, N.; Huskens, J.; Hunter, C. N.; Subramaniam, V.; Otto, C., Long-Range Energy Propagation in Nanometer Arrays of Light Harvesting Antenna Complexes, Nano Letters 2010, 10, (4), 1450–1457.

43. Tsargorodska, A.; Cartron, M. L.; Vasilev, C.; Kodali, G.; Mass, O. A.; Baumberg, J. J.; Dutton, P. L.; Hunter, C. N.; Torma, P.; Leggett, G. J., Strong Coupling of Localized Surface Plasmons to Excitons in Light-Harvesting Complexes, Nano Letters 2016, 16, (11), 6850–6856.

44. Ciesielski, P. N.; Faulkner, C. J.; Irwin, M. T.; Gregory, J. M.; Tolk, N. H.; Cliffel, D. E.; Jennings, G. K., Enhanced Photocurrent Production by Photosystem I Multilayer Assemblies, Advanced Functional Materials 2010, 20, (23), 4048–4054.

45. Ravi, S. K.; Yu, Z. M.; Swainsbury, D. J. K.; Ouyang, J. Y.; Jones, M. R.; Tan, S. C., Enhanced Output from Biohybrid Photoelectrochemical Transparent Tandem Cells Integrating Photosynthetic Proteins Genetically Modified for Expanded Solar Energy Harvesting, Advanced Energy Materials 2017, 7, (7), 1601821.

