## Supporting Information for "Proteoliposomes as energy transferring nanomaterials: enhancing the spectral range of light-harvesting proteins using lipid-linked chromophores"

### Materials and Methods

##### LHCII protein purification

Trimeric LHCII complexes from spinach were biochemically purified as described previously (Adams et al., 2018). Briefly, spinach leaves (purchased from a local supermarket) were macerated in ice-cold buffer A (300 mM sucrose, 5 mM EDTA, 50 mM HEPES, pH 7.5), the liquid filtered through muslin cloth and chloroplasts collected by centrifugation, resuspended in buffer B (5 mM EDTA, 10 mM Tricine pH 7.4) and osmotically lyzed by adding an equal volume of buffer C (400 mM sucrose, 5 mM EDTA, 10 mM Tricine, pH 7.4). Isolated thylakoid membranes were adjusted to 0.5 mg Chl/mL and then solubilized with 0.5% (w/v) detergent *n*-dodecyl α-*D*-maltoside (α-DDM, Generon) in 20 mM HEPES buffer for 1 hr on ice. Thylakoid membrane proteins were then separated via ultracentrifugation on sucrose density gradients (8-13% w/w sucrose at 100,000 x *g*, 36 hr, 4 °C). The LHCII trimer band was then collected, concentrated using 30 kDa Amicon Ultra centrifugal filters (Merck Millipore, UK), and further purified using high-resolution size exclusion chromatography in 150 mM NaCl, 0.03% α-DDM, 20 mM HEPES (pH 7.5) using a 16/600 Superdex 200 prep grade column on an AKTA Prime FPLC system (GE Healthcare Life Sciences, PA, USA). After a final concentration stage, LHCII trimers were at a concentration of approx. 100 nM in buffer of 20 mM HEPES (pH 7.5) and estimated 0.3% α-DDM. SDS-PAGE and Native-PAGE was used to confirm protein purity and oligomerisation state, as previously. Purified LHCII trimers were characterised by absorption spectroscopy at each step and before every proteoliposome preparation. LHCII trimer molar concentration is estimated as [Chl mM concentration] x 42, where "chlorophyll mM concentration" is determined from absorbance measurements after methanol/acetone pigment extraction as in (Porra et al., 1989). This leads to an approximate conversion factor from optical density (at 675nm) to LHCII trimer mM concentration has a numerical value of 5.417 x $10^{-7}$ abs/mM.

#### Reconstitution of LHCII into proteoliposomes

Plant thylakoid lipids monogalactosyldiacylglycerol (MGDG), digalactosyldiacylglycerol (DGDG), sulphoquinovosyldiacylglycerol (SQDG) and L-α-phosphatidylglycerol (Soy PG), and the synthetic lipid 1,2-dioleoyl-sn-glycero-3-phosphocholine (DOPC) were purchased from Avanti Polar Lipids as lyophilized solids. The fluorescently-tagged lipid Texas Red 1,2-dihexadecanoyl-*sn*-glycero-3-phosphoethanolamine (TR-DHPE) was purchased as a solid from Life Technologies (Invitrogen). Lipid mixtures were prepared by solubilising dry lipids with a 2:1 chloroform: methanol and mixing to obtain the desired ratios and final mass and subsequently dried under dry nitrogen gas flow for 40 minutes and then placed in vacuum desiccator overnight to remove any residual trances of chloroform (room temperature, in the dark). Lipid aliquots were then either used immediately or stored under argon gas at -80 °C until use. Our standard thylakoid lipid mixture used for all samples contained 35% MGDG, 20% DGDG, 12% SQDG, 8% Soy PG and 25% DOPC (% wt/wt), adapted from Grab and co-authors (2016). TR-DHPE was added as required to aliquots containing the standard lipid mixture before drying. Glass vials were used throughout when working with lipids in organic solvents.

Aliquots of dry thylakoid lipid mixture (as prepared above) were solubilised with 0.5% α-DDM, 20 mM HEPES (pH 7.5) at room temperature for approx. 16 hr with agitation via a pinwheel rotator to generate a mixed micellar lipid-DDM solution (approx. 9:1 molar ratio of detergent-to-lipid). The starting protein-lipid-detergent suspension was prepared in plastic microfuge tubes by mixing calculated volumes of the following: lipid-DDM solution, aqueous buffers, and purified LHCII trimers to a final concentration of: 1 mM total lipid, 0.2% α-DDM, 20 mM HEPES (pH 7.5), 40 mM NaCl and the desired LHCII concentration. The desired LHCII concentration is achieved by calculating the volume of isolated LHCII trimers required to reach a defined lipid-to-protein (mol/mol) ratio for each sample (with molar concentration of lipids calculated from known masses and molecular weights and LHCII protein concentration determined from absorption as stated above). The lipid-DDM-protein mixture was then incubated with Bio-Beads (Bio-Rad) to gradually remove the detergent and allow proteoliposome formation via self-assembly, as follows: four incubation cycles with increasing quantities of fresh Bio-Beads (8 mg/mL, 20 mg/mL and 40 mg/mL and 100 mg/mL) for 90 min, 90 min, 90 min, and ~16 hr, respectively. Proteoliposomes were prepared in sets of 5 to 7, stored in the dark at 4 °C when not in use, and diluted samples from these immediately were characterised by ensemble spectroscopies (within 16 hr) and by microscopies (within 24-72 hr).

#### Ensemble absorption and fluorescence spectroscopy (cuvette-based)

Before spectroscopy measurements, proteoliposome samples were diluted in a buffer of 40 mM NaCl, 20 mM HEPES (pH 7.5), to obtain a large enough volume for use in a 10 x10 mm quartz cuvette at a low enough absorbance of ≤0.1 at 675 nm to avoid inner filter effects (Yuan and Walt, 1957). Cuvette-based absorption spectroscopy was performed using an Agilent Technologies Cary 5000 UV-Vis-NIR absorption spectrophotometer equipped with an integrating sphere to remove any minor scattering effects (Diffuse Reflectance Accessory, Agilent).

Cuvette-based steady-state and time-resolved fluorescence spectroscopy was performed using an Edinburgh Instruments FLS980 fluorescence spectrophotometer equipped with dual excitation monochromators and dual emission monochromators. Samples were maintained at 20 °C and gently stirred during all measurements using a thermoelectrically-cooled cuvette-holder with magnetic stirring capabilities (Quantum Northwest TC 1 Temperature Controller). For steady-state emission spectra, a 450W Xenon arc lamp was used for excitation and a red-sensitive-PMT for detection (Hamamatsu R928 PMT). Emission scans with selective excitation of LHCII were acquired with excitation at 473 nm collecting emission from 500-800nm (2nm and 1nm bandwidth excitation and emission slits, respectively). Emission scans with selective excitation of Texas Red were acquired with excitation at 540 nm collecting emission from 550-800nm (1nm bandwidth for both excitation and emission slits). Data acquisition parameters were 0.5 nm steps, integrating 0.1 s/step and five scans averaged for all.

Fluorescence lifetime measurements used a 473nm pulsed diode laser or a 566nm pulsed LED for LHCII or Texas Red selective excitation, respectively. A dedicated high-speed red-sensitive PMT was used for detection (Hamamatsu H10720-20 PMT). A built-in neutral density (ND) filter wheel was applied to the pulsed laser for LHCII lifetime measurements to set excitation power as desired, approximately 37.5nW for LHCII and 4.5nW for Texas Red. Control measurements for excitation power versus fluorescence lifetime showed that singlet-singlet annihilation effects are likely to be avoided using these settings (see **supplementary Figure S11**). Decay curves from the Edinburgh FLS980 system were fitting using the manufacturer's supplied software. All ensemble spectroscopy data was further analysed in Origin Pro (v.9) graphing software.

All decay curves for TR (as in main text Figure 2C) were acquired using an alternative system, the Horiba Quantamaster fluorescence spectrometer equipped with a higher power excitation laser, due to the low Texas Red signal observed in time-resolved measurements on the most quenched samples when using the Edinburgh FL980 spectrometer. Note that this was done to improve the signal-to-noise, and we confirmed that the trends were consistent between the Edinburgh and Quantamaster systems (see **supplementary Figure S12**).

#### Epifluorescence microscopy analysis and photo-bleaching experiments

Substrates were glass coverslips 50 x 25 mm (#1.5 thickness), prepared by piranha cleaning for 40 min and used within 48 hr. Hydrophobic ultrathin adhesive imaging spacers (0.12-mm depth, 9-mm diameter) were then attached to substrates to create small wells to confine a droplet of buffer (Electron Microscopy Sciences, Hatfield, PA), for an open sample setup to allow multiple buffer exchanges. Proteoliposome samples were diluted 1/50 in a buffer of 10 mM MES (pH 6.0) 150 mM NaCl buffer and incubated with clean glass for 30 minutes in the dark (note, a lower pH buffer was used here during membrane adsorption in an attempt to promote interactions with the highly electronegative glass). Samples were then washed with seven changes of 10 mM MES (pH 6.0) 150 mM NaCl buffer in order to remove any loosely associated membranes, before being washed a final time with three changes of 20 mM HEPES (pH 7.5) 20mM NaCl to keep imaging conditions consistent with spectroscopy, returning the pH to our standard 7.5.

Epifluorescence microscopy was performed using a Nikon E600 microscope equipped with a Andor Zyla 4.2 sCMOS detector and appropriate filter cubes (LHCII cube: excitation 450-475 nm, dichroic 500 nm, emission 650-800 nm; Texas Red cube: excitation 540-580 nm, dichroic 595, emission 600-660 nm). Images were taken with a using an x40 air objective (N.A. 0.6), 1s exposure and appropriate ND filters inserted to maintain the maximum number of counts at a level for good detector signal-to-noise and linearity (10-75% of detector saturation). Two-channel imaging (Texas Red + LHCII) of a field of view was performed sequentially by switching between cubes and ND filters as appropriate.

For deliberate photo-bleaching of LHCII, an aperture was inserted to expose an approx. 30 µm diameter region of the sample for a continuously period of 120s through the LHCII filter cube at full power (i.e. no ND filters). During the bleaching, the aperture diameter was increased incrementally by approx. 10 µm every 30s to bleach multiple regions by different amounts (i.e., regions 1 to 4 in main paper **Figure 3B**). Subsequently, full-field images were acquired sequentially with LHCII and Texas Red filter cubes, to visualize the effect of photo-bleaching.

##### Fluorescence Lifetime Imaging Microscopy (FLIM) and single-proteoliposome analysis

FLIM measurements were performed on a Microtime 200 time-resolved confocal fluorescence microscope (PicoQuant GmbH). This system uses an Olympus IX73 inverted optical microscope as a sample holder with light passing into and exiting various filter units for laser scanning, emission detection and timing electronics. Excitation lasers were reflected toward the sample by a 488/561 dichroic mirror. Prior to the dichroic mirror, a small portion of the beam is deflected towards a photodiode that provides an average power readout for the excitation source before the objective lens. The excitation beam is focused through a 100X oil objective (N.A. 1.4) (UPlanSApo, Olympus). For calculating excitation power, we estimate that there is ~85% transmission efficiency at the excitation wavelengths of 485 nm and 561nm.

The excitation sources, an LDH 485 nm and an LDH 561 nm laser heads (PicoQuant), were driven in Pulsed Interleaved Excitation (PIE) mode by a PDL 828 Sepia II burst generator module (PicoQuant) at a pulse rate of 10 MHz per laser (i.e. 20 MHz overall). The pulse width for the LDH 485 nm and LDH 561 nm lasers were 90 ps and 70 ps respectively. The voltage supplied to each laser was set at the minimum required to allow lasing and kept constant for all measurements (this maintains the shortest possible pulse FWHM and provides the best temporal resolution). Laser power was set to the desired output using a combination of neutral density filters and a micro blade cut-off that partially blocks the laser beam. An excitation fluence of 0.026 mJ/cm$^2$ was used in order to limit any damage to the samples, this value was selected after trailing a series of excitation powers on control samples as shown in **Figure S13**).

Emission from the sample was passed through the same objective lens and dichroic mirror, towards a detection arm of the optical path. LHCII emission and Texas Red emission were separated by a 635LP filter in a beamsplitter tower that directed the emission towards two detectors. Emission wavelengths shorter than 635 nm, were directed though a 620/60 emission filter before being detected by a hybrid Photomultiplier Tube (PMT) detector (PicoQuant). Emission wavelengths longer than 635 nm, were directed though a 690/70 emission filter before being detected by a Single Photon Avalanche Diode (SPAD) detector (PicoQuant). With this arrangement of detectors, the PMT was optimised to detect Texas Red emission, and SPAD detected emission from LHCII. Timing electronics were a time-correlated single photon counting (TCSPC) TimeHarp 260 module (PicoQuant).

The PIE beam was directed across the sample using a “FLIMBee” mirror-based galvanometer scanner (PicoQuant). FLIM measurements were generally taken for 256 x 256 pixels across a 25 x 25 µm field of view. The dwell time for each pixel was set to 25 µs, such that an entire frame was captured in 1.64 s, and 500 frames were accumulated for each field of view. Collecting the data in this manner, allowed for the quantification of fluorophore bleaching, and the separation of frames into subgroups such that the lifetimes can be analysed at different times during the measurement. Analysis of fluorescence decay curves were performed using inbuilt fitting functions in the SymPhoTime software (PicoQuant GmbH) to fit a bi-exponential decay that was reconvoluted with measured Instrument Response Functions (IRFs). A good fit was confirmed when residuals were minimized, and Chi$^2$ was <1.1.

### Supplementary results (explanations and figures)

#### Protein biochemistry: obtaining a high purity sample of trimeric LHCII

Denaturing and native gel electrophoresis was performed on all LHCII preparations. Representative gels are shown below.

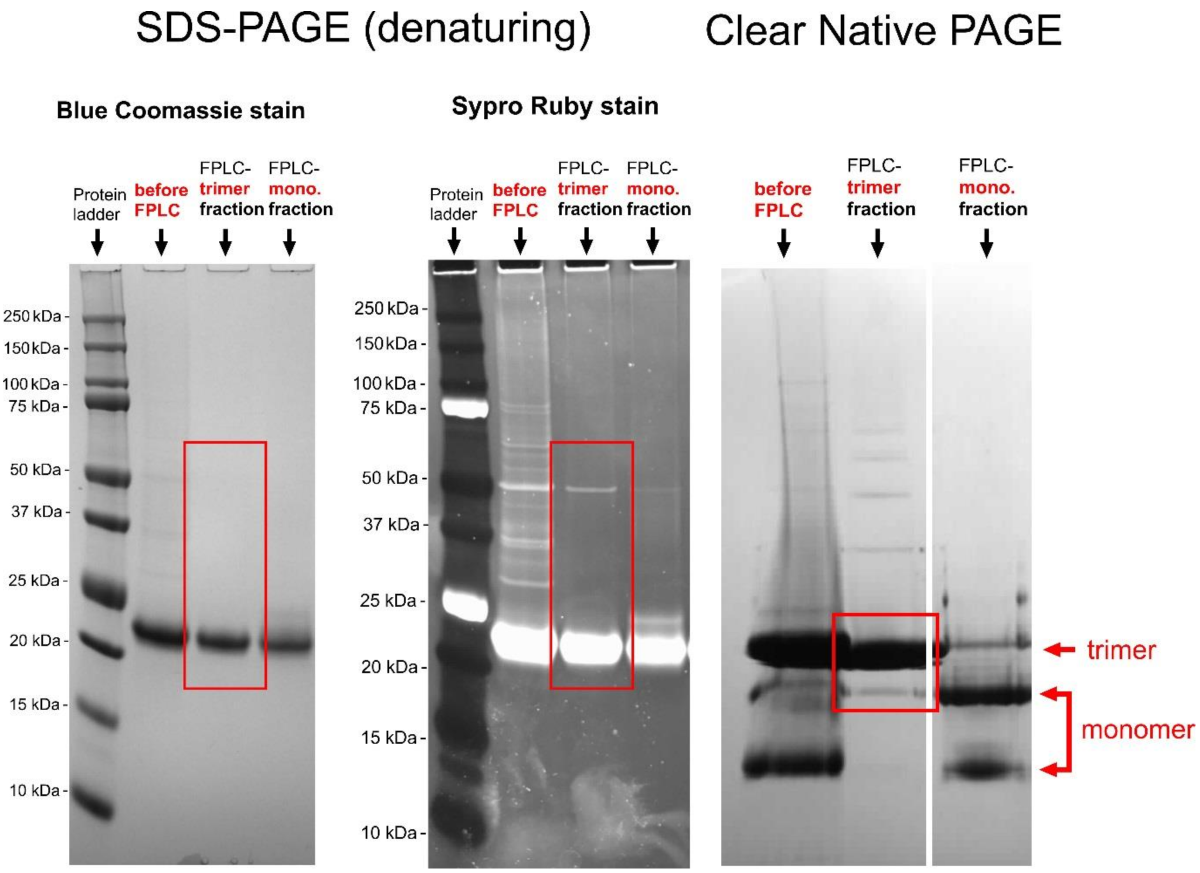


***Figure S1*** *SDS-PAGE gel with either Coomassie* (left panel) *or SYPRO-RUBY stain* (middle panel). *Channels show, in order: protein standard, LHCII with impurities (pre size exclusion chromatography), purified trimeric LHCII (post size exclusion chromatography), purified trimeric LHCII (post size exclusion chromatography). Native-PAGE gel* (right panel) *was ran at 4°C and then stained with Coomassie. Channels show, in order: LHCII with impurities (pre size exclusion chromatography), purified trimeric LHCII (post size exclusion chromatography), purified trimeric LHCII (post size exclusion chromatography).*

#### Analytical Ficoll gradient analysis of lipid and protein co-localisation

Analytical ultracentrifugation of selected proteoliposomes and control samples on Ficoll density gradients was performed to observe the differential sedimentation of any sub-populations. A standard quantity of each samples (150 uL undiluted/by mass) was loaded onto 5-20% continuous Ficoll gradients in SW55 tubes (in 20 mM HEPES, 40 mM NaCl buffer). Note, Ficoll has a similar density to sucrose but aqueous solutions have a much lower osmolality, making it a gentler medium for isolation of vesicles in which osmotic lysis is prevented. Gradients were centrifuged at 226,000 x*g* for 10 hr at 4 °C (with no brake) and then immediately photographed.

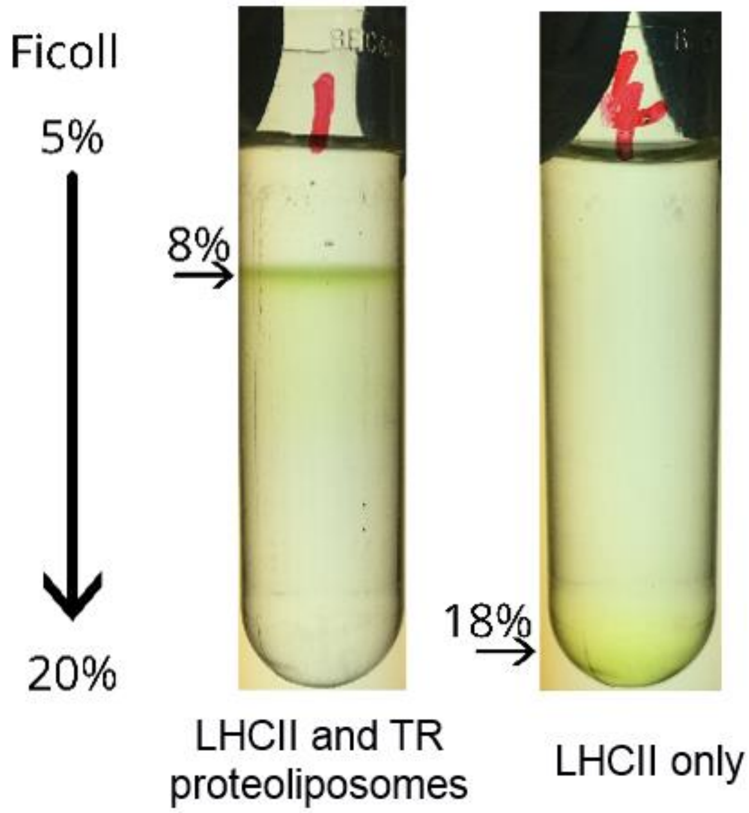


***Figure S2*** *Analytical Ficoll gradients between 5-20%* ***(Left)*** *Representative proteolispomes sample (0.7µM LHCII, 8µM Texas Red, 1mM total lipids) showing a clear single population and no aggregates of LHCII* ***(Right)*** *LHCII in detergent at a 0.7µM concentration showing clear sedimentation of LHCII not reconstituted into proteoliposomes.*

#### Observation of minimal changes to LHCII fluorescence emission spectra

Shown below are fluorescence emission spectra from proteoliposome Series 2, all normalized to an emission of 1.0 at the peak maximum around 681 nm. Shifts of wavelength (in the x-axis) are no more than 1 nm for any sample. Broadening of the peaks increases with decreasing LHCII concentration to a maximum of 19% of the peak area compared to isolated LHCII, suggest that up to 19% of chlorophylls may have an altered environment. Whilst not ideal this is entirely in line with multiple other proteoliposome studies (Adams et al., 2018; Natali et al., 2016; Pandit et al., 2011)where LHCII appears to be destabilized when the protein-to-lipid ratio is very low. We speculate that LHCII-LHCII interactions may be stabilizing. This is outside of the scope of the current work. For the current study, whilst subtle changes to spectra are detected it is important to note that the vast majority of LHCII appears undamaged and that we can still calculate accurate FRET and enhancement of LHCII emission.

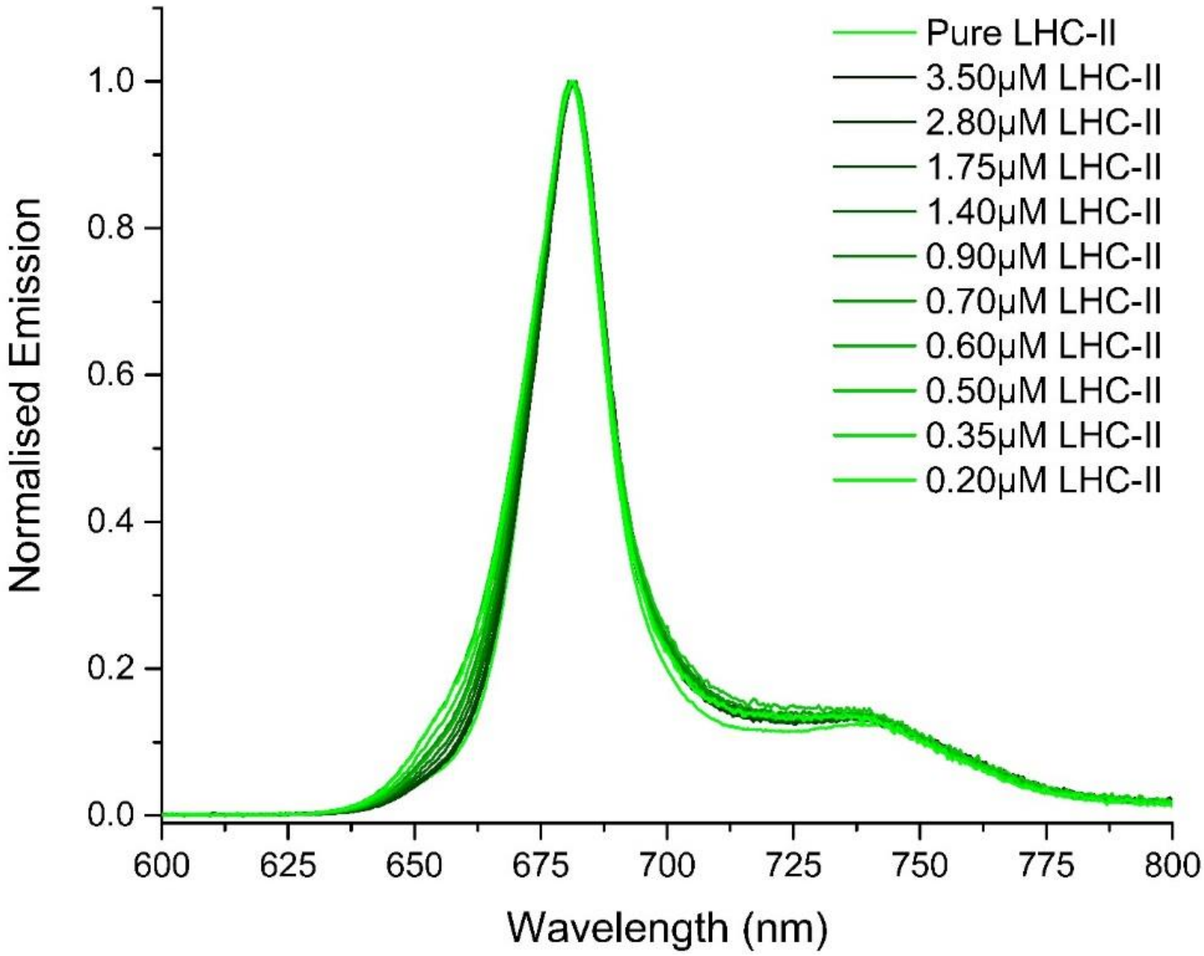


***Figure S3*** *Normalised steady-state emission spectra of sample Series 2 with selective LHCII excitation at 473nm. Note minimal emission peak shift and broadening with different LHCII concentrations. All measurements were taken in a buffer of 20mM HEPES (pH 7.5), 40mM NaCl.*

#### Quantification of the LHCII and Texas Red concentration in proteoliposomes by deconvolution analysis of the absorption spectra

All analysis, below, was performed using Origin Pro (v.9) graphing software. All absorption spectra were "baselined" to give an absorbance of zero at 800nm as expected in these samples to remove any differences in noise (e.g. detectors, slight differences in cuvettes, etc.). All absorption spectra were then corrected for dilution by multiplying by dilution factor.

LHCII content was estimated before any deconvolution because there is no significant absorption from Texas Red >630 nm. LHCII content assessed from the integrated area from 635-700 nm, with LHCII concentration then calculated as:
AREA x $1.485 \times 10^{-5}$ $mM^{-1}$ (value from absorption of a known concentration of LHCII in detergent).
Area was used to ensure that any slight peak broadening did not lead to a loss of relative absorption.

Before quantification of concentration of Texas Red, the component peaks within absorption spectra were deconvoluted because LHCII overlaps throughout the TR absorption range. To obtain the TR-only component, a representative LHCII absorption spectrum (originally collected from LHCII in detergent or in liposomes) was normalised to the test sample's absorbance at 675 nm (LHCII Chl *a* $Q_y$ peak) and then subtracted, to give a result as in **supplementary Figure S4**, below.

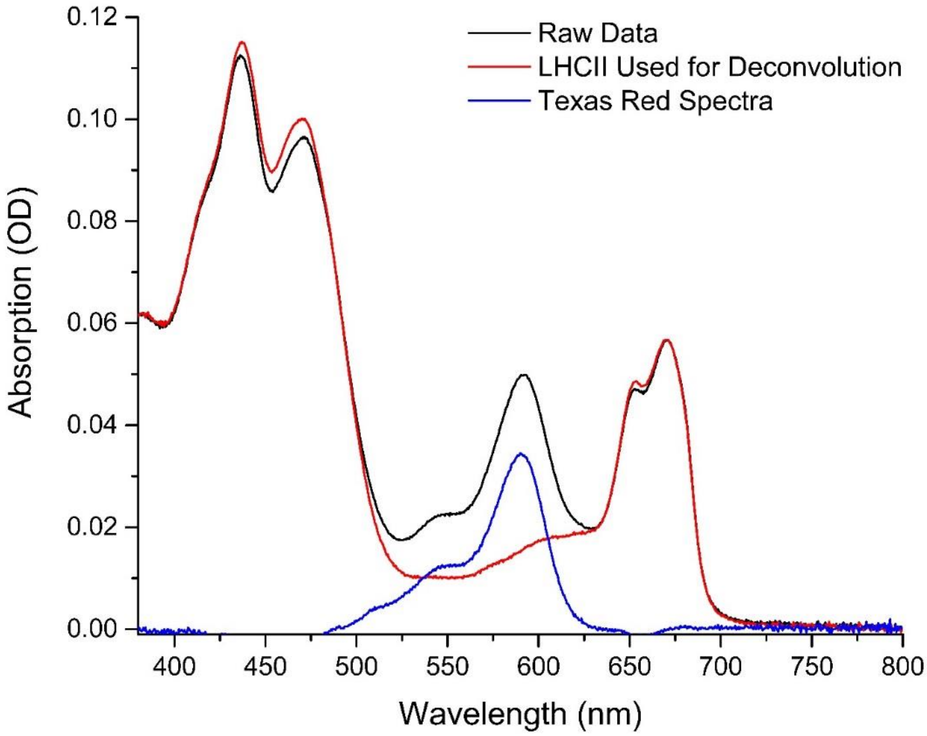


***Figure S4*** *Example of absorption spectra de-convolution for a representative proteoliposome sample (0.7µM LHCII, 12.5µM TR, 1mM total lipids). Absorption spectra from a LHCII-only control sample (*Red*) is fitted to the 675nm emission peak of the raw absorption data (*Black*) and subtracted. The resulting spectra is the absorption from only the Texas Red component in each sample (*Blue*).*

Texas Red content was assessed on these deconvoluted spectra from the peak height at 591nm with TR-DHPE concentration then calculated as:
PEAK HEIGHT x $2.09 \times 10^{-2}$ $mM^{-1}$ (value is the extinction coefficient of TR-DHPE in a 1 cm pathlength cuvette provide by the supplier in agreement with other published works).

#### Time resolved fluorescence data showing non-quenched Texas Red and LHCII

Fluorescence lifetime traces of isolated (not quenched) Texas Red (**Figure S5A**) and LHCII (**Figure S5B**) with calculated amplitude-weighted lifetimes of 4.4 ns of 3.9 ns respectively. These values were used to calculate the relative fluorescence lifetime quenching of components when reconstituted into proteolispomes, where quenching occurs due to energy transfer in the case of Texas Red and self-quenching in the case of LHCII. The instrument response function (IRF) measures the scattering of laser excitation from non-fluorescent control samples to determine the fastest possible response of the detectors. The IRF is then used when fitting decay curves and obtaining amplitude-weighted lifetimes.

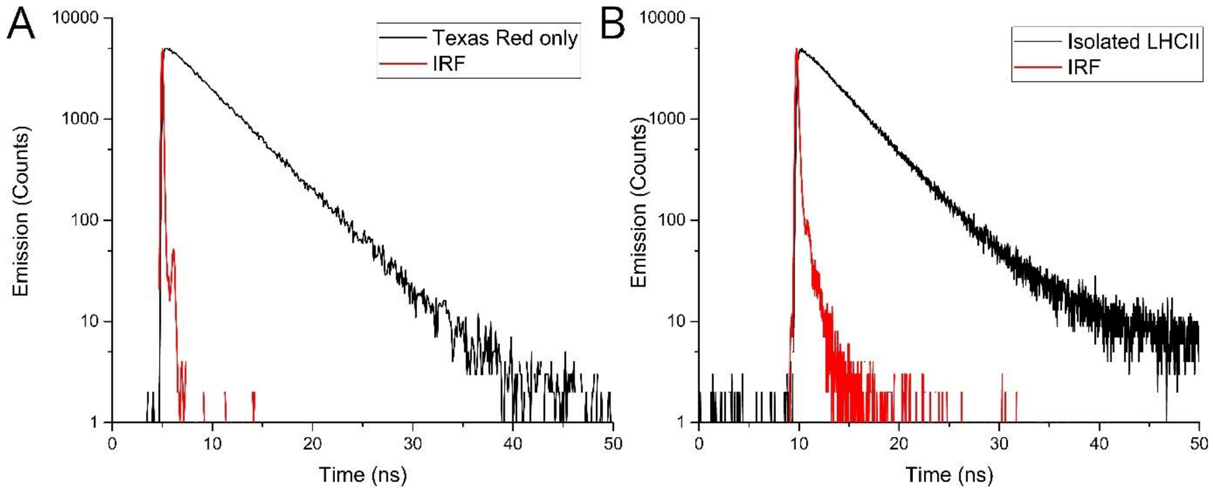


***Figure S5*** *Time resolved fluorescence data from (A) solution of liposomes comprised of 2.1 µM TR-DHPE and 1.0 mM total lipids in a buffer 10mM HEPES, pH 7.5 (absorbance ~0.1 at 561 nm). Taken using QuantaMaster fluorescence spectrometer with supercontinuum pulsed laser at 0.5MHz, excitation set to 540nm (5nm slit) and 610nm emission (1nm slit). Instrument response function (IRF) taken using colloidal silica beads. (B) LHCII trimers in a buffer of 0.03% w/v α-DDM, 10 mM HEPES, pH7.5 (absorbance ~0.1 at 675 nm). Taken using Edinburgh Instruments F980 fluorescence spectrometer with 475nm pulsed laser at 0.5MHz. Instrument response function (IRF) taken using colloidal silica beads*

#### Disruption of proteoliposomes with detergent

Shown below is the fluorescence emission spectra of a proteoliposome sample under selective Texas Red excitation (540nm) before and after solubilisation in detergent. The detergent (DDM) is known to disrupt lipid bilayers and will, after several minutes of mixing, isolate all lipids and LHCII trimers into detergent micelles. The isolation of lipids and proteins will limit any TR-to-LHCII energy transfer as they will be too spatially distant for FRET to occur and also limit any self-quenching of LHCII due to protein-protein interactions. Preventing these TR-LHCII and LHCII-LHCII interactions which cause quenching will result in regeneration of fluorescence of both components as seen in **Figure S6.** Texas Red fluorecence emission intensity increases from 3% to 87% as compared to a control sample of TR-DHPE isolated in detergent. This regeneration confirms that both LHCII and Texas Red quenching occur due to the membrane archetecure and nanoscale interactions.

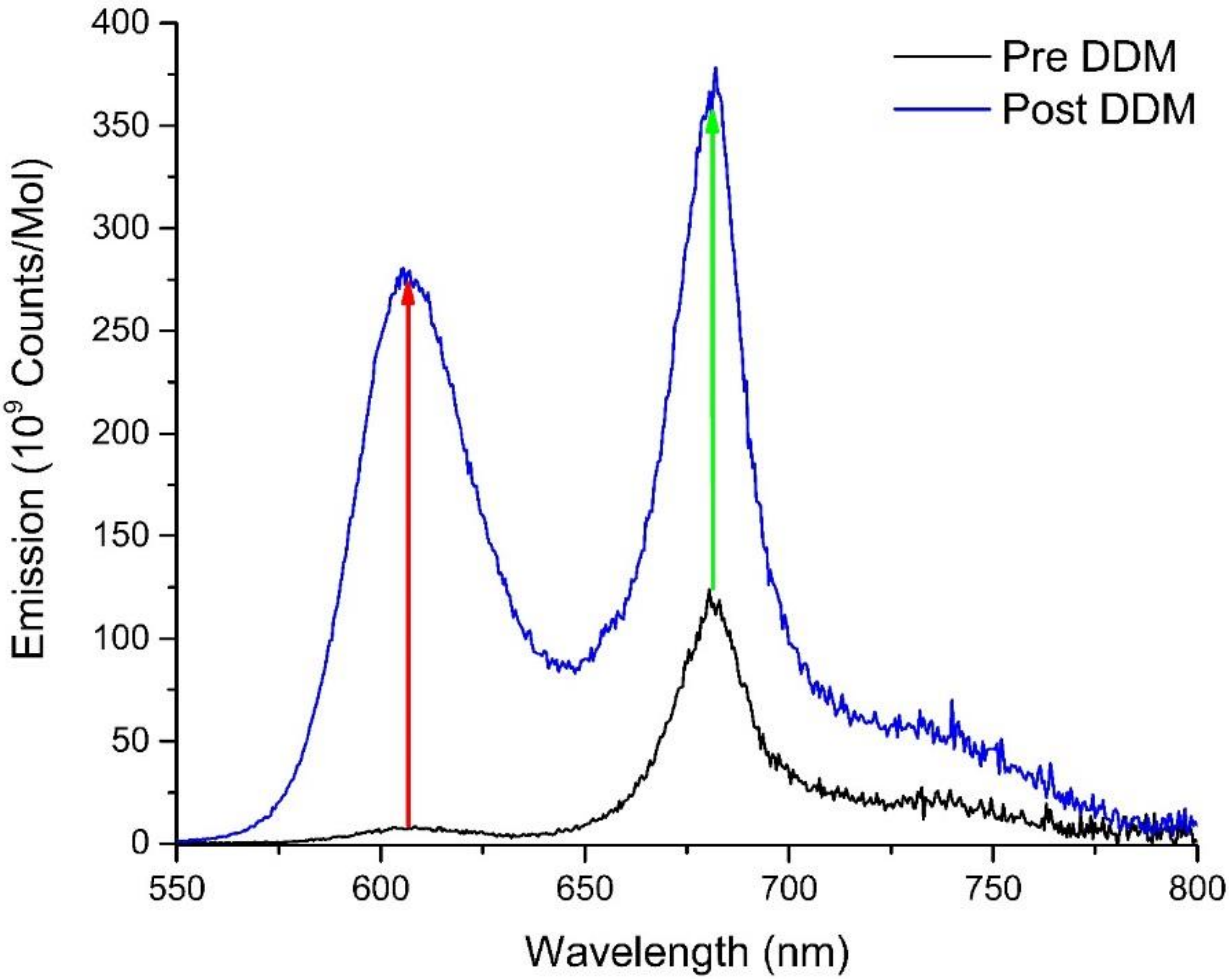


***Figure S6*** *Fluroescence emission data from 3.5µM LHCII, 12µM Texas Red, 1mM total lipid proteoliposomes diluted x60 in 20 mM HEPES 40 mM NaCl. Post DDM spectra taken after 0.1% w/w α-DDM added to cuvette and stirred for 5 minutes in order to solubilise lipids and proteins. Red and green arrows highlight the fluorescnece emission de-quenching of TR and LHCII respectivly.*

#### Quantification of the “Relative fluorescence” of LHCII and Texas Red by deconvolution analysis of fluorescence spectra

All analysis, below, was performed using Origin Pro (v.9) graphing software. Before quantification of relative fluorescence intensity data, the LHCII and Texas Red component peaks within fluorescence emission spectra were deconvoluted as in **supplementary Figure S7**, below. To obtain the LHCII-only component, a representative Texas Red emission spectrum (originally collected from Texas Red only liposomes) was normalised to the test sample’s emission at 610nm (Texas Red emission peak) and then subtracted (result as in **supplementary Figure S7A**). To obtain the Texas Red-only component, a representative LHCII emission spectrum was normalised to roughly 80% of the test sample’s emission at 681nm (result as in **supplementary Figure S7B**). This was % value was tweaked and iterated to produce a deconvoluted spectrum for TR where the value at 681nm is 17.7% of its peak maximum at 610nm, as expected for pure Texas Red. All emission spectra were then corrected for dilution by multiplying by dilution factor. It was critical that our values for emission intensity were accurate for all sample sets acquired over many weeks apart, therefore, we performed control measurements each day to check the absolute consistency of the FL980 fluorescence spectrophotometer, relative to a known isolated LHCII in α-DDM sample. Values were always within 3% of the previously acquired data.

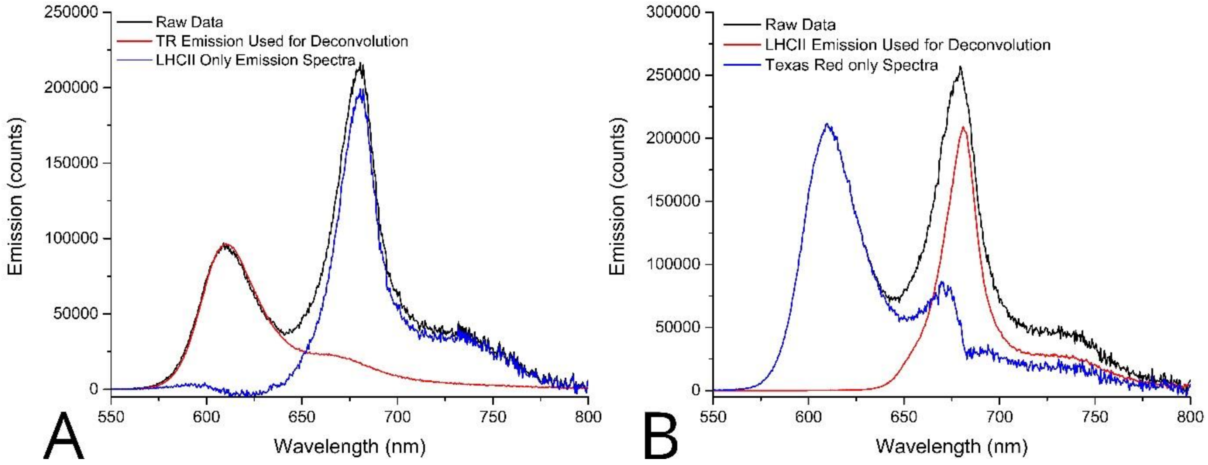


***Figure S7*** *Example of emission spectra de-convolution for a representative proteoliposome sample (0.7µM LHCII, 12.5µM TR, 1mM total lipids).* ***(A)*** *Removal of TR emission from convoluted spectra: TR only emission (red) is normalised to 610nm peak of convoluted spectra (Black) and removed leaving LHCII emission only (Blue****)*** ***(B)*** *Removal of LHCII emission from convoluted spectra: LHCII only emission (Red) is normalised to 681 peak of convoluted spectra (Black) and subtracted so that the TR emission only spectra (Blue) height at 681nm is 17.7% of height at 610nm (the same ratio as TR only emission from control sample).*

The “relative fluorescence intensity” per molecule of interest was defined as the calculated value for emission intensity divided by the molar concentration as calculated in the above section *Quantification of the LHCII and Texas Red concentration in proteoliposomes by deconvolution analysis of the absorption spectra*. Thus, “relative LHCII fluorescence intensity” (LHCII emission per mole LHCII) was calculated by integrating the spectrum between 500-800nm (post-deconvolution) and dividing this value by LHCII concentration (in mM).

And, “relative TR fluorescence intensity” (TR emission per mole TR) was calculated as the emission peak height at 610nm (post-deconvolution) and dividing this value by TR-DHPE concentration (in mM).

#### Observation of significant “self-quenching” of LHCII due to its clustering within lipid bilayers in agreement with previous studies

The know phenomenon of LHCII self-quenching when reconstituted into proteolispomes was quantified in order to accurately determine the fluorescence enhancement of LHCII due to energy transfer from Texas Red. Steady-state and time-resolved fluorescence spectra were measured on samples of LHCII both isolated in α-DDM detergent micelles and when reconstituted into proteolispomes at a comparable concentration to main text proteoliposome series 1 (0.7 µM LHCII). Steady-state fluorescence emission data (**Figure S8A**) shows a 59% decrease in LHCII emission when reconstituted into proteoliposomes compared to isolated LHCII in detergent (from $2.51 \times 10^9$ counts/mol to $1.06 \times 10^9$ counts/mol). A similar decrease of 49% is observed in LHCII fluorescence lifetime from 3.96 ns for LHCII isolated in detergent to 2.02 ns when reconstituted into proteoliposomes (**Figure S8B**). “Enhancement” of LHCII fluorescence emission with increasing Texas Red concentration in proteoliposome series 2 (main text Figure 2D) were calculated relative to this expected value for LHCII emission for proteoliposomes containing 0.7 µM LHCII, in order to take any quenching due to LHCII-LHCII interactions into account.

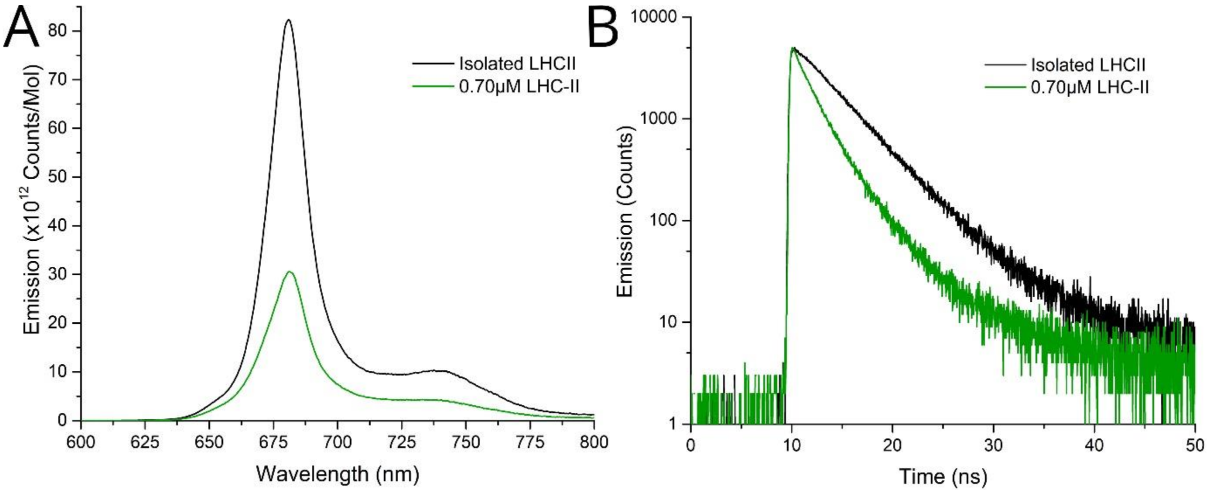


***Figure S8*** *Quantification of LHCII self-quenching when reconstituted into proteoliposomes (0.7 µM LHCII and 1mM total lipids, in a buffer of 20 mM HEPES 40 mM NaCl) vs isolated (in a buffer of 20 mM HEPES, 40 mM NaCl, 0.03% α-DDM)* ***(A)*** *Steady-state fluorescence emission data taken with selective LHCII excitation (473nm)* ***(B)*** *Time-resolved fluorescence emission data taken using Edinburgh instruments F980 fluorescence spectrometer with 475nm pulsed laser at 0.5MHz.*

#### Observation of a small degree of “self-quenching” of Texas Red in liposomes

Steady-state and time-resolved fluorescence emission spectra of Texas Red were measured for samples omitting LHCII in order to determine a value for fully emissive TR (no quenching) which could be used to calculate relative TR quenching and therefore energy transfer efficiency. TR-DHPE was measured when isolated in chloroform and α-DDM detergent micelles in addition to when reconstituted into liposomes at a comparable concentration to in sample series 2. Each sample of isolated TR-DHPE was measured at different dilutions and the final results averaged (n=3). Three individually prepared samples of TR-DHPE in liposomes were measured and the results averaged. Slight differences in the magnitude of TR fluorescence emission ‘per mol’ and amplitude-weighted lifetime were measured (**Figure S9A** and **S9B,** respectively): in detergent 8.95x$10^{11}$ counts/mol and 5.09 ns, in chloroform 8.79x$10^{11}$ counts/mol and 4.14 ns, in liposomes 8.55 x$10^{11}$ counts/mol and 4.21 ns. Differences in fluorescence may be attributed to different local solvent environments, supported by the shifts to emission peaks in the x-direction. Self-quenching of TR-DHPE in liposomes cannot be ruled out, but is not considered further in this study. These values for steady-state and time-resolved fluorescence of Texas Red in liposomes were used as the non-quenched value for energy transfer efficiency calculations as it is most representative of the local environment of Texas Red in samples, i.e., $F_D$ and $\tau_D$ in eqn. 1 and 2 for main text Figure 2D.

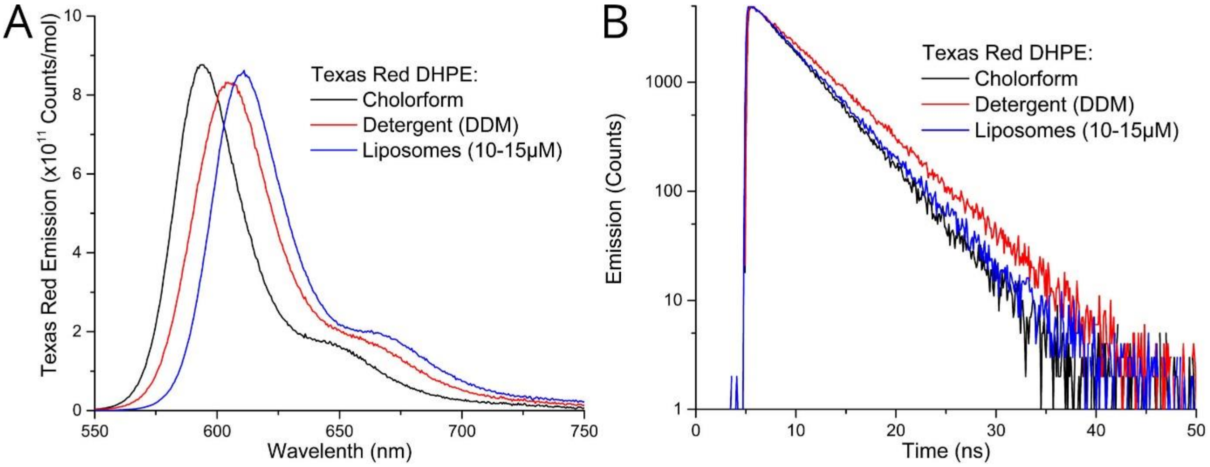


***Figure S9*** *Quantification of TR-DHPE fluorescence when in chloroform, detergent micelles and incorporated into liposomes, all samples omitting LHCII. Chloroform used was analytical grade and 99.9% pure, detergent samples used a buffer of 20 mM HEPES 40 mM NaCl 0.1% α-DDM, all liposomes samples were at a total lipid concentration of 1mM and TR-DHPE between 10-15 μM in a buffer of a buffer of 20 mM HEPES 40 mM* ***(A)*** *Steady-state fluorescence emission of Texas Red ‘per mol’ in different solvent conditions each spectra an average of three samples. Taken with selective Texas Red excitation at 540nm* ***(B)*** *Time-resolved fluorescence emission of Texas Red in different solvent conditions. Representative decay curves from one sample per condition. Taken using QuantaMaster fluorescence spectrometer with supercontinuum pulsed laser at 0.5MHz, excitation set to 540nm (5nm slit) and 610nm emission (1nm slit).*

#### Display of multiple fields from all three tested proteoliposomes/ liposomes

In **supplementary figure S10**, we present representative (25 X 25 µm) fields of view for proteoliposomes containing a "high" concentration of LHCII (2.8 µM LHCII, 12 µM TR, 1mM total lipids), a "low" concentration of LHCII (1.2µM LHCII, 12 µM TR, 1 mM total lipids), and a control "TR-only" liposome sample containing TR-DHPE without any proteins (12 µM TR, 1 mM total lipids).

In the TR-only liposome sample, there is little/no signal in the LHCII channel, and the signal observed can be attributed to the spectral overlap of the emission filters and detectors (quantifed overlap of <1% of the TR signal). In the TR channel, the donor lifetime is non-quenched, and is represented by longer "red" lifetimes in false-colour mapping of fluorescence lifetimes.

In the proteoliposome samples, we observe a clear LHCII signal for both samples (99% confidence that these signals are above the TR overlap and detector noise thresholds), with a high degree of colocalisation with the corresponding TR signals (as discussed in the main text). The presence of LHCII in both of these samples correspond to the quenching of the TR which can be observed qualitatively as the preponderance of shorter "bluer" lifetimes in the FLIM false-colour mapping (as compared to the TR-only sample).

The sample with high LHCII concentration appears on-average to show shorter "bluer" TR lifetimes and lower TR intensities than the low LHCII concentration sample when comparing across multiple fields of view; therefore providing an initial indication of the higher extent of donor quenching in the prescence of a higher acceptor concentration. However, the variety of proteoliposomes and corresponding lifetimes within a field of view presents the need for a comprehensive single-particle anaylsis.

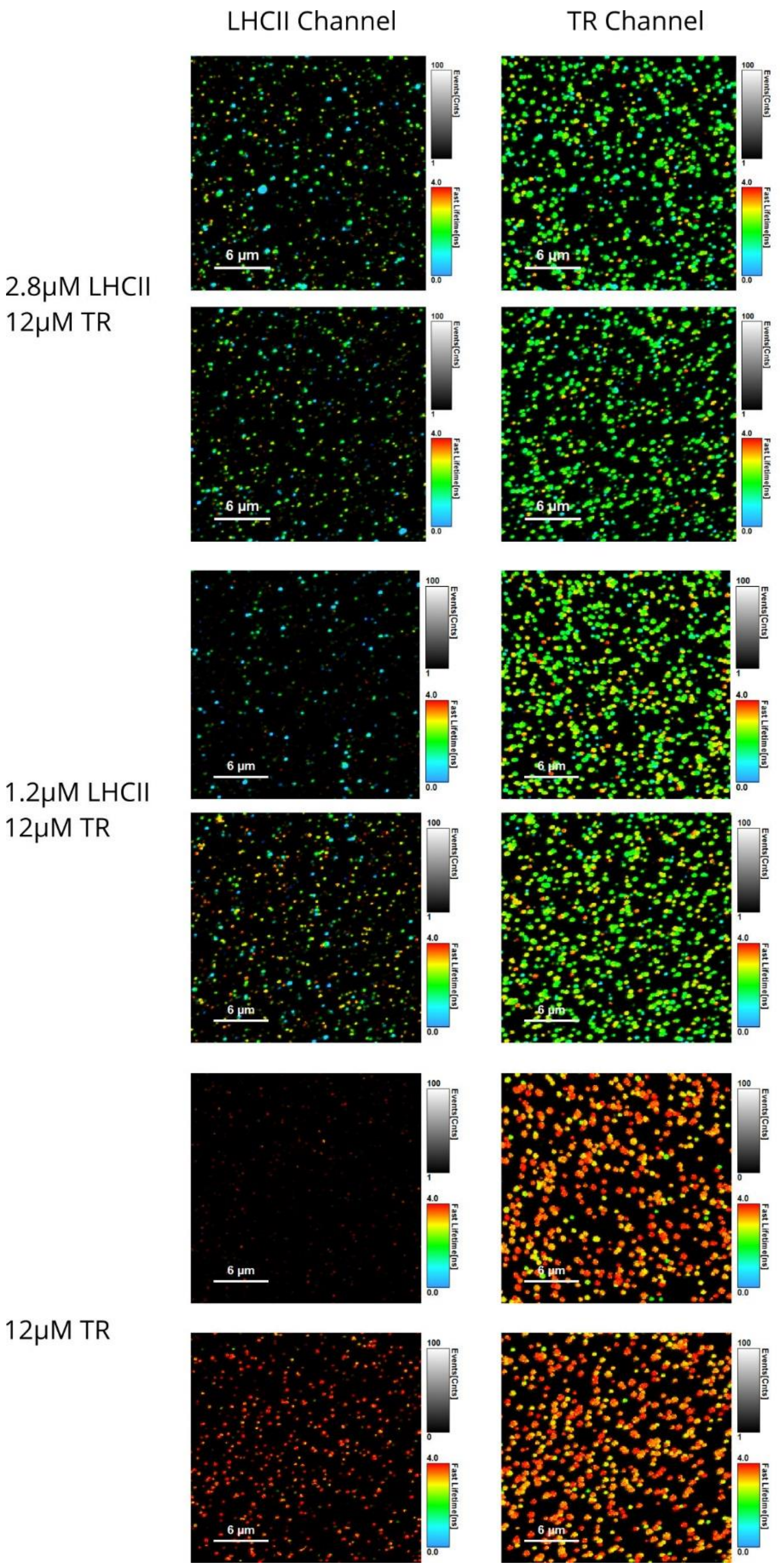


***Figure S10*** *Gallery of FLIM images.*

#### Optimization of the ensemble time-resolved fluorescence acquisitions: excitation power

A series of excitation powers were trialled on control samples of isolated LHCII, to analyze how fluorescence lifetime varies with laser excitation power and determine a “safe” low power which will not adversely damage the sample. For example, it is known that high excitation fluences can cause singlet-singlet annihilation events in LHCII aggregates which reduce the LHCII lifetime (Barzda et al., 2001). In **supplementary Figure S11**, we observed that LHCII mean lifetime was relatively constant at 3.75 to 3.85 ns for low powers of 0.1 – 1 µW, but started to decrease at power higher than 1 µW. To avoid such singlet-singlet annihilation effects (decrease in measured lifetime at high laser power), all subsequent LHCII lifetimes were measured using laser power at under 5% of this threshold. For Texas Red lifetimes, see the following page.

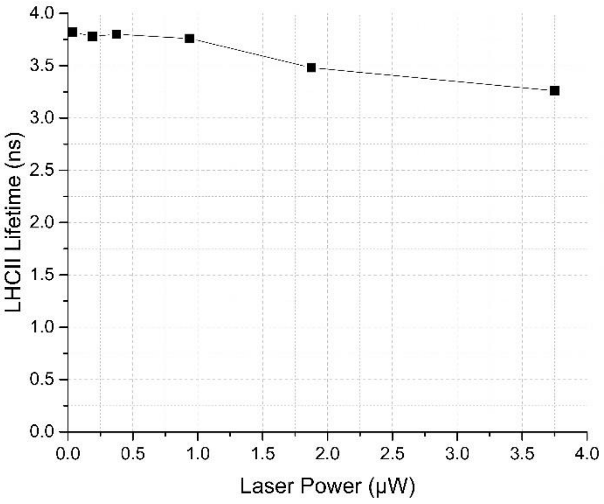


***Figure S11.*** *Measured fluorescent lifetime of isolated LHCII (20 mM HEPES, pH 7.5, 0.03% α-DDM) taken using Edinburgh instruments F980 fluorescent spectrometer at a range of laser powers. Left and right displays the full y-range of 0-4 ns or a zoomed in range (for clarity).*

#### Agreement of time-resolved fluorescence data from our standard FL-980 fluorescence spectrophotometer and a higher power Quantamaster system

Due to the low Texas Red signal observed in time-resolved measurements on the most quenched samples when using the Edinburgh FL980 spectrometer, the data for TR decay curves (as in main text Figure 2C) was acquired using a Horiba Quantamaster fluorescence spectrometer equipped with a higher power excitation laser. Decay curves from the Quantamaster system were fitting using DecayFit Fluorescence Decay Analysis software v 1.3. The Texas Red lifetime values obtained from both spectrometers were shown to be consistent between the Edinburgh and Quantamaster instruments (**supplementary Figure S12**) over the series of proteoliposome samples, however, the Quantamaster providing decay curves data with an improved higher signal to noise. Therefore, data from the Quanmaster system is used for main text Figure 2C and 2D. Furthermore, the agreement between the magnitude of TR fluorescence lifetime where laser power is very low (LED in our standard FL-980 system) or moderate (supercontinuum laser in the Quantamaster system) gives us confidence that are no significant annihilation effects with Texas Red. This is agreement with the finding that Texas Red lifetime does not change with laser fluence in FLIM (see next page).

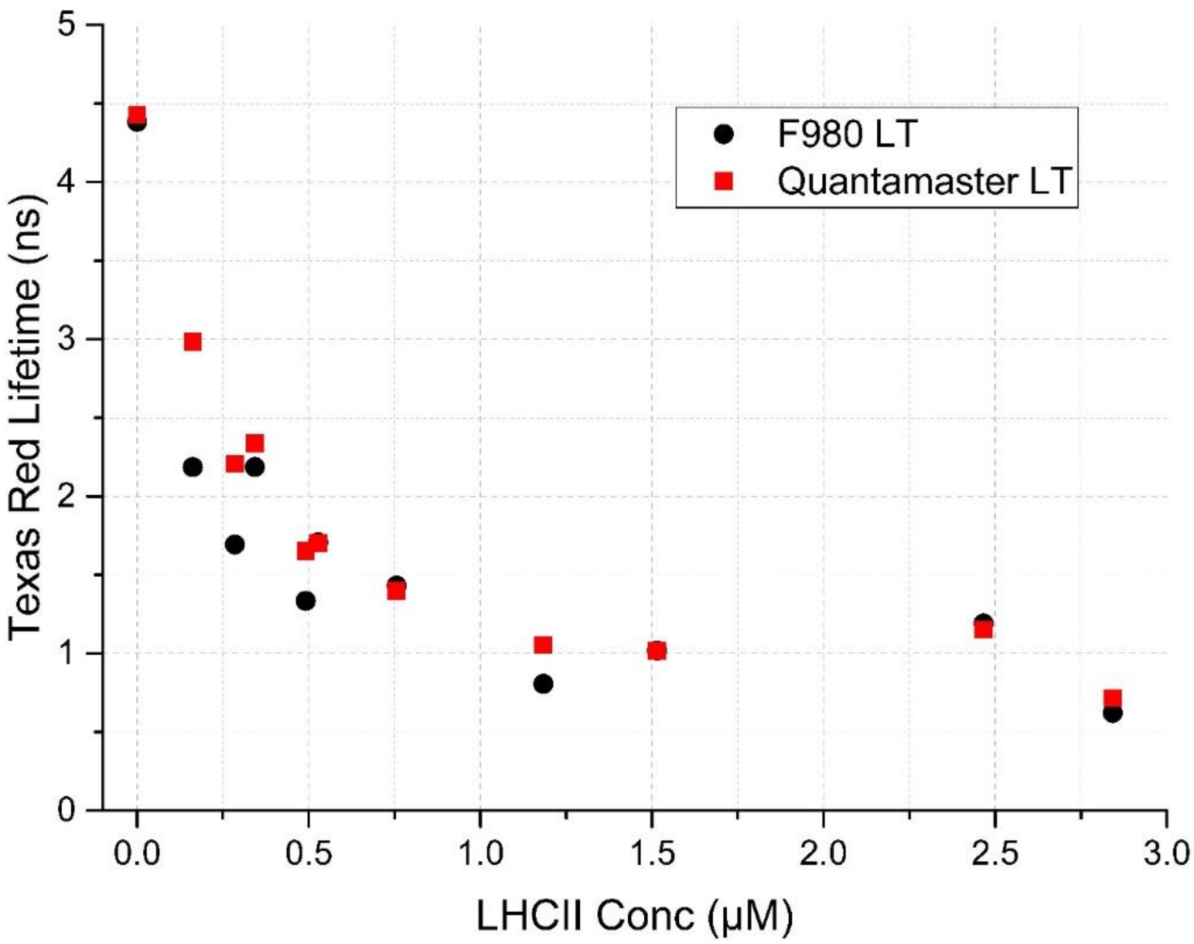


***Figure S12*** *Measured TR fluorescncent lifetimes from sample series 1 (Constant TR concentration, varying LHCII concentration) taken using Edinburgh Instruments F980 spectrometer (Black datapoints) and Horiba Quantamaser spectrometer (Red datapoints). The lifetimes recorded on both systems are very similar suggesting that no high power annihilation is occuring when using the higher power Quantamaster system.*

#### Optimization of the FLIM acquisitions: excitation power

A series of excitation powers on the FLIM were trialled on control samples of either isolated LHCII (in detergent solution, laser focussed into the droplet) and Texas Red at low concentration in liposomes (absorbed to the glass substrate as usual). Excitation fluence (power per unit area) at the sample surface were calculated to be from 0.002 mJ/cm$^2$ to 0.388 mJ/cm$^2$. As for the cuvette-based measurements (above), control measurements were performed for FLIM to analyze how fluorescence lifetime varies with laser excitation power and determine a “safe” low power which will not adversely damage the sample.

In **supplementary Figure S13**, we observed that LHCII mean lifetime was 3.5 to 4.0 ns for low fluences of 0.002 to 0.026 mJ/cm$^2$ but started to decrease at fluence higher than 0.026 mJ/cm$^2$ and was <1 ns at 0.388 mJ/cm$^2$. In contrast, TR mean lifetime was observed at all tested laser powers, so one assumes that fluence was well below the threshold of annihilation effects.

Therefore, in all FLIM acquisitions reported the current study, an excitation fluence of 0.026 mJ/cm$^2$ was used for both lasers, because it provided a balance between a high enough power to produce reasonable signal but low enough that neither LHCII nor TR undergo significant annihilation events.

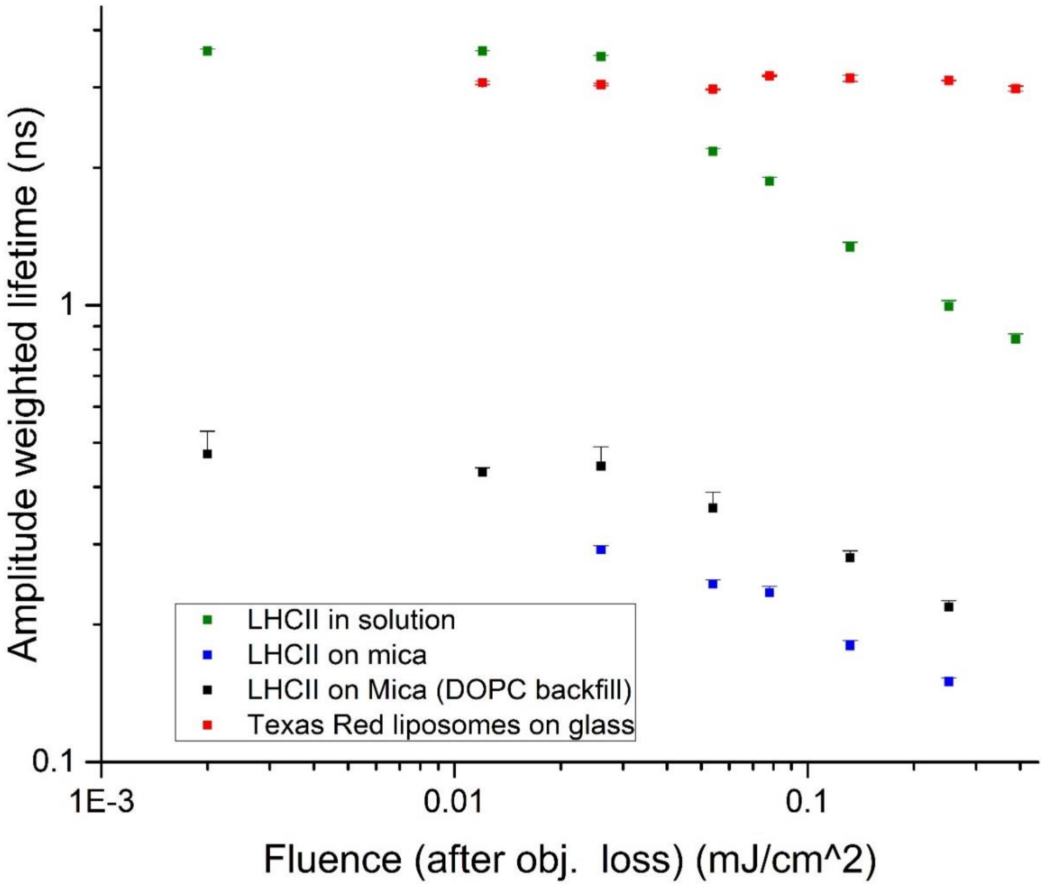


***Figure S13*** *Measured fluorescent lifetimes of isolated LHCII and TR liposomes taken using FLIM system at a range of laser fluences.* Green datapoints*: emission from isolated LHCII in α-DDM solution with variation in the 488 nm laser power.* Red datapoints*: emission from Texas Red in liposomes with variation in the 561 nm laser power.*

#### Ensemble spectroscopy tabulated data

**Table S1** below shows the ensemble spectroscopy data used to calculated LHCII emission enhancement in proteoliposome series 1 and energy transfer efficiency in proteoliposome series 2. LHCII absorbance is measured as the integrated area between 635-800 nm from sample absorption spectra and then used to calculate the concentration of LHCII in liposomes (μM) as stated in **Figure S4.** The weight/weight fraction of LHCII in liposomes taking into account lipid mass and concentration is also calculated. Texas Red absorbance is measured as the optical density of the Texas Red absorption peak from the deconvoluted absorption spectra and used to calculate concentration in liposomes as stated in **Figure S4.** The molar fraction of TR-DHPE compared to total lipids is also calculated.

Steady-state and time-resolved fluorescence of Texas Red are measured with selective excitation, with steady-state spectra deconvoluted as in **Figure S7.** Relative TR Emission (steady-state fluorescence intensity) and Calculated TR $<\tau>$ (mean lifetime) are used to estimate the energy transfer efficiency (ETE) with eqn. 1 and 2 as in main text **Figure 2.** Relative LHCII emission (steady-state fluorescence intensity) is measured with selective Texas Red excitation in order to determine the enhancement due to Texas Red to LHCII energy transfer, from spectra deconvoluted as in **Figure S7.** LHCII Emission Enhancement is a measure of the % increase of Relative LHCII Emission as compared to LHCII liposomes containing a comparable LHCII concentration to that in proteolispomes series 1 (as described in **Figure S8**).

| Sample description | | Measured LHCII Absorbance | Calculated LHCII Concentration | | Measured TR Absorbance | Calculated TR Concentraion | | Relative TR Emission per mol TR | Calculated TR <T > | ETE (steady-state) | ETE (time-resolved) | Relative LHCII Emission per mol LHCII | LHCII Emission Enhancement |
|---|---|---|---|---|---|---|---|---|---|---|---|---|---|
| | | Integranted area (635-800nm) | (μM) | (wt/wt% LHCII/total) | OD at 591nm | (μM) | (mol/mol lipid%) | (540nm Excitation) | (ns) | (%) | (%) | (540nm Excitation) | (% Enhancemnt relative to 0.7μM LHCII sample with no TR) |
| LHCII-0.70μM/ | No TR | 2.33 | 0.52 | 7.4 | 0.000 | 0.00 | 0.00 | - | - | - | - | 1.03E+08 | - |
| LHCII-0.70μM/ | TR-2.5μM | 2.36 | 0.53 | 7.5 | 0.005 | 1.65 | 0.17 | - | - | - | - | 1.33E+08 | 25 |
| LHCII-0.70μM/ | TR-5.0μM | 2.56 | 0.57 | 8.1 | 0.012 | 3.90 | 0.39 | - | - | - | - | 1.57E+08 | 50 |
| LHCII-0.70μM/ | TR-7.5μM | 2.46 | 0.55 | 7.8 | 0.021 | 6.54 | 0.65 | - | - | - | - | 1.76E+08 | 83 |
| LHCII-0.70μM/ | TR-10.0μM | 2.50 | 0.56 | 7.9 | 0.027 | 8.35 | 0.84 | - | - | - | - | 1.92E+08 | 106 |
| LHCII-0.70μM/ | TR-12.5μM | 2.42 | 0.54 | 7.7 | 0.034 | 10.76 | 1.08 | - | - | - | - | 2.10E+08 | 129 |
| LHCII-0.70μM/ | TR-15.0μM | 2.50 | 0.56 | 7.9 | 0.040 | 12.53 | 1.25 | - | - | - | - | 2.34E+08 | 153 |
| LHCII-0.70μM/ | TR-17.5μM | 2.57 | 0.57 | 8.1 | 0.048 | 15.09 | 1.51 | - | - | - | - | 2.47E+08 | 181 |
| LHCII-0.70μM/ | TR-20.0μM | 2.53 | 0.56 | 8.0 | 0.054 | 16.88 | 1.69 | - | - | - | - | 2.67E+08 | 171 |
| LHCII-0.70μM/ | TR-22.5μM | 2.45 | 0.54 | 7.7 | 0.064 | 19.97 | 2.00 | - | - | - | - | 2.93E+08 | 221 |
| LHCII-0.70μM/ | TR-25.0μM | 2.73 | 0.61 | 8.6 | 0.070 | 21.92 | 2.19 | - | - | - | - | 2.96E+08 | 214 |
| No LHCII/ | TR-12.0μM | 0.00 | 0.00 | 0.0 | 0.024 | 7.61 | 0.76 | 1.06E+12 | 4.43 | N/A | N/A | - | - |
| LHCII-0.20μM/ | TR-12.0μM | 0.73 | 0.16 | 2.4 | 0.047 | 14.86 | 1.49 | 3.77E+11 | 2.98 | 55.9 | 29.2 | - | - |
| LHCII-0.35μM/ | TR-12.0μM | 1.28 | 0.28 | 4.2 | 0.044 | 13.72 | 1.37 | 2.31E+11 | 2.21 | 73.0 | 47.5 | - | - |
| LHCII-0.50μM/ | TR-12.0μM | 1.54 | 0.34 | 5.0 | 0.030 | 9.48 | 0.95 | 2.52E+11 | 2.34 | 70.6 | 44.4 | - | - |
| LHCII-0.60μM/ | TR-12.0μM | 2.21 | 0.49 | 7.0 | 0.041 | 12.94 | 1.29 | 1.43E+11 | 1.65 | 83.3 | 60.8 | - | - |
| LHCII-0.70μM/ | TR-12.0μM | 2.37 | 0.53 | 7.5 | 0.028 | 8.88 | 0.89 | 1.54E+11 | 1.70 | 82.0 | 59.6 | - | - |
| LHCII-0.90μM/ | TR-12.0μM | 3.40 | 0.76 | 10.4 | 0.029 | 8.97 | 0.90 | 1.13E+11 | 1.40 | 86.8 | 66.7 | - | - |
| LHCII-1.40μM/ | TR-12.0μM | 2.66 | 1.18 | 15.4 | 0.022 | 14.07 | 1.41 | 4.95E+10 | 1.05 | 94.2 | 75.1 | - | - |
| LHCII-2.75μM/ | TR-12.0μM | 3.40 | 1.51 | 18.9 | 0.016 | 9.75 | 0.97 | 5.51E+10 | 1.02 | 93.6 | 75.8 | - | - |
| LHCII-2.80μM/ | TR-12.0μM | 3.69 | 2.47 | 27.5 | 0.010 | 9.38 | 0.94 | 3.68E+10 | 1.15 | 95.7 | 72.7 | - | - |
| LHCII-3.50μM/ | TR-12.0μM | 3.19 | 2.84 | 30.5 | 0.011 | 14.27 | 1.43 | 2.07E+10 | 0.72 | 97.6 | 82.9 | - | - |

***Table S1*** *Display of data shown in main text Figure 2 for sample series 1 and 2 (top and bottom respectively). Raw data values are shown in black, calculated values shown in red.*

#### Single particle FLIM tabulated data

In order to calculate the fluorescence lifetimes on a per-proteoliposome basis, manual analysis was performed by fitting fluorescence decay curves extracted from a region-of-interest defined to represent individual proteoliposomes. Only well-resolved proteoliposomes with sufficient signal to produce a good fit were selected (criteria of counts >500 and fit chi$^2$ <1.2). This selection process may produce inherent bias as we do not assess the weakly emitting particles, but nevertheless the trends are clear.

**Table S2** below shows representative single-proteoliposome analysis of FLIM data which is used, firstly, to estimate TR and LHCII co-localization (comparison of their intensities) and, secondly, to produce the population distribution (frequency histograms) in main text **Figure 4C**. Individual particles were selected and numbered for consistency and to avoid repeat measurements. A perimeter was then drawn around each particle in order to select a defined number of pixels considered to make up that individual particle. The intensities of LHCII and Texas Red signal from each particle were measured and detector bleed-through was corrected for. "Corrected signal" is after subtraction for spectral overlap in the opposite channel (Raw LHCII signal contains significant bleedthrough of TR fluorescence, so this was quantified with control measurements and removed; in contrast there is minimal bleedthrough of LHCII fluorescence into the TR channel). The amplitude-weighted fluorescence lifetime of Texas Red ($T_{average}$) was then calculated from a bi-exponential decay function fitted to the produced fluorescence decay curves. $A_1$, $T_1$ and $A_2$, $T_2$ represent the amplitude ($A$) and lifetime ($T$) of the two exponential components. The chi$^2$ fit quality parameter is shown (chi$^2$ <1.2 represent a good fit). The lifetime of LHCII is not assessed in the current study.

| Particle number | Pixels in partice | LHCII Raw Signal | LHCII corrected Signal | TR Raw Signal | TR Corrected Signal | TR $A_1$ | TR $A_2$ | TR $\tau_1$ | TR $\tau_2$ | TR $\tau_{average}$ | TR $\chi^2$ |
|---|---|---|---|---|---|---|---|---|---|---|---|
| # | # | Counts | Counts | Counts | Counts | Counts | Counts | ns | ns | ns | |
| 1 | 42 | 204 | 102 | 10572 | 10561 | 560 | 355 | 2.17 | 0.00 | 1.33 | 0.96 |
| 2 | 32 | 88 | 47 | 3903 | 3897 | 61 | 20 | 0.76 | 2.70 | 1.22 | 0.96 |
| 3 | 21 | 284 | 259 | 2364 | 2351 | 24 | -87 | 2.30 | 0.03 | 0.84 | 1.01 |
| 4 | 39 | 40 | -10 | 4760 | 4756 | 894 | 243 | 2.44 | 0.00 | 1.90 | 0.97 |
| 5 | 30 | 111 | 78 | 3047 | 3041 | 102 | 96 | 2.70 | 0.74 | 1.76 | 1.04 |
| 6 | 20 | 115 | 100 | 1287 | 1281 | 180 | 17 | 0.93 | 3.00 | 1.20 | 0.71 |
| 7 | 23 | 67 | 43 | 2255 | 2251 | 155 | 345 | 3.20 | 1.30 | 1.91 | 1.17 |
| 8 | 34 | 40 | 15 | 2053 | 2049 | 74 | 6 | 1.40 | 4.70 | 1.60 | 0.76 |
| 9 | 63 | 563 | 430 | 13605 | 13578 | 570 | 1000 | 2.50 | 0.77 | 1.40 | 1.01 |
| 10 | 57 | 1557 | 1484 | 7054 | 6986 | 288 | 1200 | 2.40 | 0.67 | 1.00 | 1.06 |

***Table S2*** *Representative single-particle analysis of histogram data shown in main text Figure 4.*

#### Estimation of energy transfer efficiency in single proteoliposomes via FLIM

We know from our photobleaching experiments that a quenching of LHCII leads to an increase in TR emission (so-called donor "de-quenching") and we know that this occurred over our FLIM measurements. We attempted to correct for this effect to provide a more reliable estimate of what the TR lifetime would be, for the calculation of FRET efficiency (our main interest). Single-proteoliposome data of TR mean lifetime were corrected to remove the effect of LHCII bleaching as follows. The TR mean lifetime averaged over the entire field of proteoliposomes within an image was assessed for a period of the initial 170 s of an acquisition where LHCII bleaching was minimal (frames 1 – 50), compared to the full period of acquisition (frames 1 – 500). From this the TR mean lifetime change over the entire image was determined to be: a 53% increase for the high-LHCII proteoliposomes and a 52% increase for low-LHCII proteoliposomes, as shown in **supplementary Table S3**, below.

Single-proteoliposome analysis was performed on the full 1-500 frames of data, necessary to provide a large enough photon count level for an adequate decay curve for a good fit (typically 3000 counts with a range from 1000-10000). The data without any correction applied is shown in main text **Figure 4C**. These raw values for TR mean lifetimes were then adjusted by the average relative change in lifetime determined as above (52 and 52%) to give a representation of the mean TR lifetime unbiased by photobleaching effects. Energy transfer efficiency (ETE) was then calculated using the conventional relationship corrected TR lifetimes, $ETE_{LT} = 1 - \frac{\tau_{DA}}{\tau_D}$ for the high-LHCII proteoliposomes and the low-LHCII proteoliposomes (as *DA*) using the TR-only liposome sample FLIM data as the donor only sample (as *D*). This FRET efficiency estimation is shown in main text **Figure 4D**.

| | High LHCII | | Low LHCII | |
|---|---|---|---|---|
| Frames | Caclulated TR <T> | Error | Caclulated TR <T> | Error |
| # | ns | ns | ns | ns |
| 1-50 | 0.713 | 0.018 | 0.712 | 0.026 |
| 1-500 | 1.335 | 0.008 | 1.360 | 0.007 |
| Ratio | 0.534 | - | 0.524 | - |

***Supplementary Table S3.*** *The difference in measured TR lifetime over frames 1-50 vs 1-500.*

#### References


Adams, P.G., Vasilev, C., Hunter, C.N., and Johnson, M.P. (2018). Correlated fluorescence quenching and topographic mapping of Light-Harvesting Complex II within surface-assembled aggregates and lipid bilayers. Biochimica et Biophysica Acta, Bioenergetics *1859*, 1075-1085.

Barzda, V., Gulbinas, V., Kananavicius, R., Cervinskas, V., van Amerongen, H., van Grondelle, R., and Valkunas, L. (2001). Singlet–Singlet Annihilation Kinetics in Aggregates and Trimers of LHCII. Biophysical Journal *80*, 2409-2421.

Grab, O., Abacilar, M., Daus, F., Geyer, A., and Steinem, C. (2016). 3D-Membrane Stacks on Supported Membranes Composed of Diatom Lipids Induced by Long-Chain Polyamines. Langmuir *32*, 10144-10152.

Natali, A., Gruber, J.M., Dietzel, L., Stuart, M.C., van Grondelle, R., and Croce, R. (2016). Light-harvesting Complexes (LHCs) Cluster Spontaneously in Membrane Environment Leading to Shortening of Their Excited State Lifetimes. Journal of Biological Chemistry *291*, 16730-16739.

Pandit, A., Shirzad-Wasei, N., Wlodarczyk, L.M., van Roon, H., Boekema, E.J., Dekker, J.P., and de Grip, W.J. (2011). Assembly of the major light-harvesting complex II in lipid nanodiscs. Biophysical Journal *101*, 2507-2515.

Porra, R.J., Thompson, W.A., and Kriedemann, P.E. (1989). Determination of accurate extinction coefficients and simultaneous equations for assaying chlorophylls a and b extracted with four different solvents: verification of the concentration of chlorophyll standards by atomic absorption spectroscopy. Biochimica et Biophysica Acta, Bioenergetics *975*, 384-394.

Yuan, P., and Walt, D. (1957). Calculation for Fluorescence Modulation by Absorbing Species and Its Application to Measurements Using Optical Fibers. Analytical Chemistry *59*, 2391-2394.
