## Supplementary figures and images for "Proteoliposomes as energy transferring nanomaterials: enhancing the spectral range of light-harvesting proteins using lipid-linked chromophores"

### Video 1 - LHCII

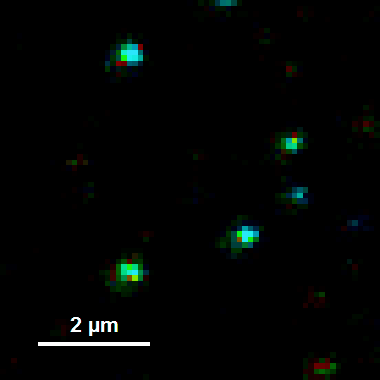

### Video 1 - TR

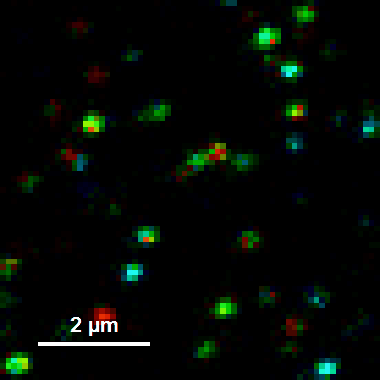
